# The Australasian dingo archetype: *De novo* chromosome-length genome assembly, DNA methylome, and cranial morphology

**DOI:** 10.1101/2023.01.26.525801

**Authors:** J. William O. Ballard, Matt A. Field, Richard J. Edwards, Laura A.B. Wilson, Loukas G. Koungoulos, Benjamin D. Rosen, Barry Chernoff, Olga Dudchenko, Arina Omer, Jens Keilwagen, Ksenia Skvortsova, Ozren Bogdanovic, Eva Chan, Robert Zammit, Vanessa Hayes, Erez Lieberman Aiden

**Author notes:** **Correspondence address.**. School of Biosciences, University of Melbourne, Royal Parade, Parkville, Victoria 3052, Australia.

## Abstract

**Background:** One difficulty in testing the hypothesis that the Australasian dingo is a functional intermediate between wild wolves and domesticated breed dogs is that there is no reference specimen. Here we link a high-quality *de novo* long read chromosomal assembly with epigenetic footprints and morphology to describe the Alpine dingo female named Cooinda. It was critical to establish an Alpine dingo reference because this ecotype occurs throughout coastal eastern Australia where the first drawings and descriptions were completed.

**Findings:** We generated a high-quality chromosome-level reference genome assembly (Canfam_ADS) using a combination of Pacific Bioscience, Oxford Nanopore, 10X Genomics, Bionano, and Hi-C technologies. Compared to the previously published Desert dingo assembly, there are large structural rearrangements on Chromosomes 11, 16, 25 and 26. Phylogenetic analyses of chromosomal data from Cooinda the Alpine dingo and nine previously published *de novo* canine assemblies show dingoes are monophyletic and basal to domestic dogs. Network analyses show that the mtDNA genome clusters within the southeastern lineage, as expected for an Alpine dingo. Comparison of regulatory regions identified two differentially methylated regions within glucagon receptor GCGR and histone deacetylase HDAC4 genes that are unmethylated in the Alpine dingo genome but hypermethylated in the Desert dingo. Morphological data, comprising geometric morphometric assessment of cranial morphology place dingo Cooinda within population-level variation for Alpine dingoes. Magnetic resonance imaging of brain tissue show she had a larger cranial capacity than a similar-sized domestic dog.

**Conclusions:** These combined data support the hypothesis that the dingo Cooinda fits the spectrum of genetic and morphological characteristics typical of the Alpine ecotype. We propose that she be considered the archetype specimen for future research investigating the evolutionary history, morphology, physiology, and ecology of dingoes. The female has been taxidermically prepared and is now at the Australian Museum, Sydney.

## Introduction

The most influential book on evolution, Darwin’s 1859 *On the origin of species* [1], starts with a chapter on domestication to reverse engineer natural selection. Some nine years later Darwin [2] expanded his initial thinking into the book *The variation of animals and plants under domestication*. He hypothesized that the process of domestication proceeded in a stepwise manner first by unconscious selection (wild →tamed) followed by what we now call artificial selection (tamed →domesticated), with the key distinction between these processes being the involvement of humans on mating and reproduction. A gap in our ability to test Darwin’s hypothesis has been the identification of a model system with an extant plant or animal that is intermediate between the wild ancestor and the domesticate. Here we explore the overarching hypothesis that the Australasian dingo (*Canis (familiaris) dingo*) is evolutionarily intermediate between the wild wolf (*Canis lupus*) and domestic dogs (*Canis familiaris*) [3]. One alternate hypothesis is that the process of domestication is continual and does not proceed in a stepwise manner [4], instead representing a series of phases reflecting an intensification of the relationship between a wild animal (or plant) and human societies [5].

The taxonomic name of the dingo remains unstable, however, it is now clear the Australasian dingo is a distinct evolutionary lineage closely related to domestic dogs [6]. The first European drawing of an animal referred to as a “dingo” appears in White 1790 [7] with a more complete anatomical description appearing in Meyer 1793 [8]. A “large dog” from coastal eastern Australia near Sydney was earlier illustrated by George Stubbs in 1772, based on a recorded description by Joseph Banks from 1770; it is now clear that this animal was a dingo, but the name had not yet been learned from the local Aboriginal people. We follow the precedent that when zoologists disagree over whether a certain population is a subspecies or a full species, the species name may be written in parentheses. Scientists advocating a General Lineage Species Concept consider dingoes to be distinct species (*Canis dingo*) or a subspecies of domestic dog (*Canis familiaris dingo*) [9-11]. Others advocating a Biological Species Concept [12] consider the dingo to be a breed of dog (*Canis familiaris* breed dingo) due to the interfertility between dingo and domestic dog [11, 13, 14].

Corbett [15] mentioned the possibility of three different dingo ecotypes existing in north, central and southeastern Australia. These are now referred to Tropical, Desert, and Alpine dingoes [16]. Subsequently, Corbett [17] noted that dingo skulls from southeastern Australia (Alpine dingoes) were genuinely different from those of the rest of the country, but posited the differences may be due to hybridization with domestic dogs rather than independent lineages. Jones [18] agreed that the southeastern dingoes, were distinct and suggested a revaluation of ecotype morphologies to resolve the conundrum.

Analyses of mitochondrial variation in canids from Southeast Asia supports the hypothesis that there are distinct dingo lineages [19-22]. Zhang et al. [19] found a strong Bayesian posterior value supporting the separation of Australian dingoes into two groups. One is a northwestern group, whereas the other is a southeastern group that clusters with New Guinea Singing dogs (*Canis (familiaris) hallstromi*). Support for two, or perhaps three, distinct lineages of dingoes has also come from Y-chromosome and SNP-chip data [23, 24].

The dog is the first species to be domesticated [25]. They are likely the most frequently kept domestic animal, exhibit exceptional levels of morphological variation, and many breeds have been developed by strong artificial selection in the past 200 years [26-28]. The Australasian dingo has been proposed to be a functional [29] and evolutionary [6] intermediate between wild wolves and domesticated dogs. Unfortunately, the absence of a dingo holotype reference specimen impedes our ability to definitively determine whether dingoes are a tamed intermediate or a feral canid because we do not have a single reference point that links the scientific name to a specific specimen [30].

This study aims to link high resolution long-read *de novo* chromosomal assembly, mitochondrial DNA sequence and the DNA methylome with morphological descriptions of head shape and computed tomography data of brain data to describe the ‘archetype’ dingo (Fig. 1). This designation will support future comparisons with a reference enabling further characterization of the evolutionary history of the dingo. In this case we do not propose any formal taxonomic name for the specimen as it is a regional morphotype that is being characterized however we suggest the principle of having a ‘type’ specimen makes biological sense and will enable the focusing of future research.

**Figure 1.**
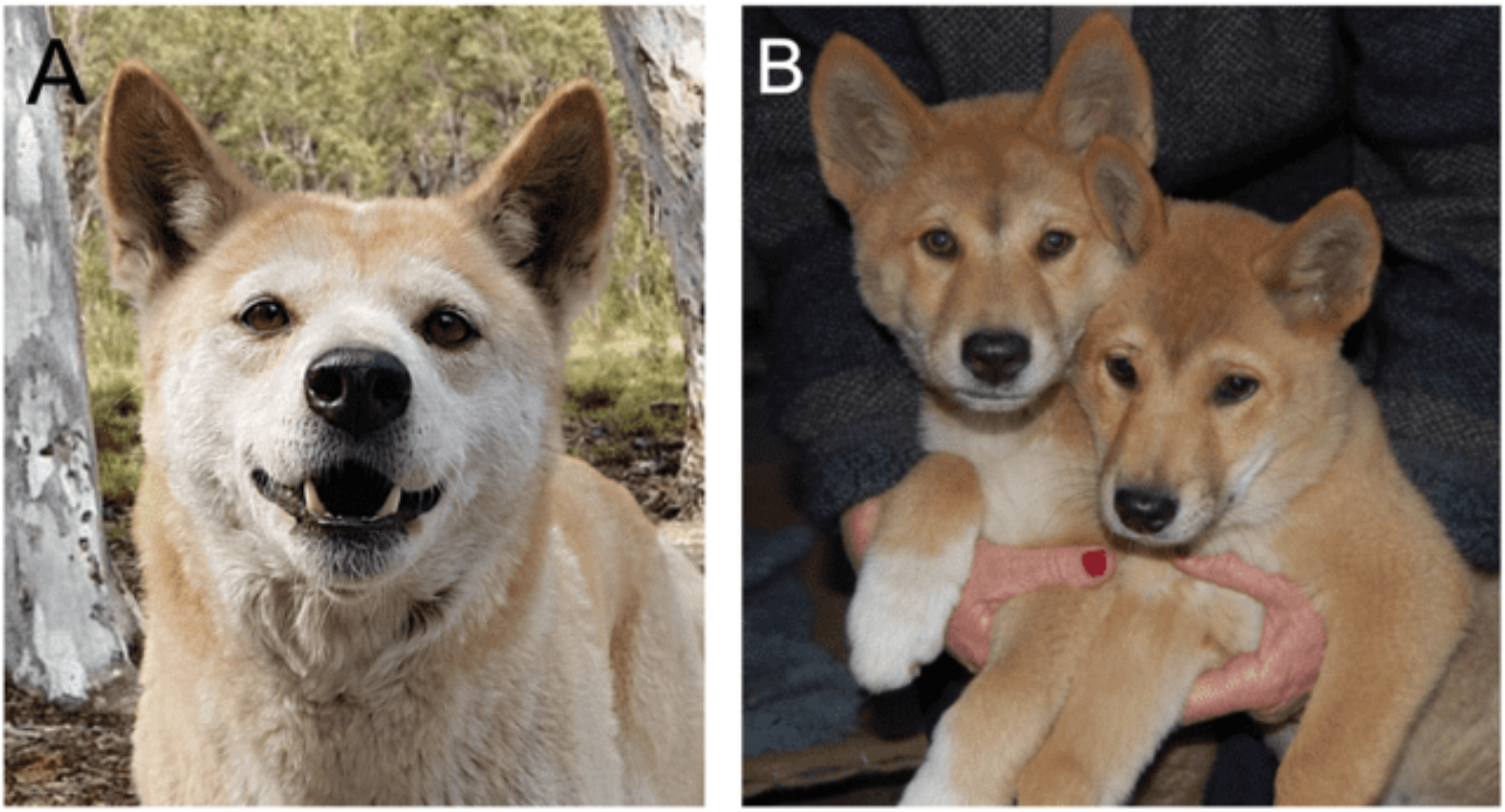
Cooinda the dingo. The genomic and morphological data in this study is based upon a single individual named Cooinda from Dingo Sanctuary Bargo in the southern highland region of New South Wales. Based on her parentage, broad skull, and stocky appearance the Sanctuary considers her an Alpine dingo. We compare her with other dingoes found in southeastern Australia and with those found in the center and northwest of the continent including Desert dingo Sandy [6]. (A) Dingo Cooinda as an adult female. (B) Brother Typia (RHS) and Cooinda (LHS) as 8-week-old puppies.

## Results

### Chromosome-level genome assembly

#### Workflow

The genome was assembled following a similar pipeline to Field et al. [28] (Supplementary Fig. 1). Briefly, 1722 contigs were assembled from SMRT and ONT sequence data with a total length of 2.38 Gb and N50 length of 12.4 Mb [31]. The contig assembly was then polished for two rounds with SMRT reads, correcting ∼5 million bases in the first round and ∼15 thousand in the second [32, 33]. The assembled sequence contigs were scaffolded sequentially using 10X linked-reads and polished with 10X linked-reads [33]. The scaffolded assembly was then super scaffolded with Bionano and Hi-C proximity ligation.

Supplementary Fig. 2 shows the contact matrices generated by aligning the Hi-C data set to the genome assembly after Hi-C scaffolding [34, 35]. To increase the contiguity of the assembly we used the SMRT and ONT reads to fill gaps, which was then followed by a final round of SMRT read polishing. The gap filling successfully closed 282 gaps increasing contig N50 to the final figure of 23.1 Mb. A final round of polishing was performed with 10X linked reads. The resulting chromosome-length genome assembly and its gene annotation was deposited to NCBI with accession number GCA_012295265.2.

#### Assembly statistics and completeness

The final assembly had a total length of 2,398,209,015 bp in 477 scaffolds with a scaffold and contig N50 of 64.8 Mb and 23.1 Mb, respectively (Table 1). Chromosome-level scaffolds accounted for 98.4 % of the assembly with only 0.9 % (21.1 Mb) of all sequences not aligning to a CanFam4.1 chromosome [36].

**Table 1:** Genome assembly and annotation statistics for Alpine dingo (Cooinda) vs Desert dingo assembly (Sandy)

| Statistic | Alpine dingo | Desert dingo |
| --- | --- | --- |
| Total sequence length | 2,398,209,015 | 2,349,862,946 |
| Total ungapped length | 2,390,794,485 | 2,349,829,267 |
| Number of contigs | 802 | 228 |
| Contig N50 | 23,108,747 | 40,716,615 |
| Contig L50 | 36 | 20 |
| Number of scaffolds | 477 | 159 |
| Scaffold N50 | 64,752,584 | 64,250,934 |
| Scaffold L50 | 15 | 14 |
| Number of gaps | 325 | 69 |
| BUSCO complete (single/<br>duplicate copy) | 95.1% (S: 92.7% D:2.4%) | 95.3% (S: 92.9% D:2.5%) |
| BUSCO fragmented | 0.8% | 0.8% |
| BUSCO missing | 4.1% | 3.8% |

Evaluation by Benchmarking Universal Single-Copy Orthologs (BUSCO v5.2.2 [37]) against Carnivora_odb10 data set (n=14,502) indicated that 95.1 % of the conserved single-copy genes were complete (Table 1, Supplementary Fig. 3A). Only 3 of 13,791 complete (single-copy or duplicated) BUSCO genes were not on the 39 nuclear chromosome scaffolds.

Next, we compared single-copy “Complete” BUSCO genes in Alpine dingo Cooinda and nine canid genomes [6, 27, 28, 36, 38-41]). Of the 13,722 genes, 13,711 were found in the assembly using BUSCOMP v1.0.1. Only Sandy the Desert Dingo v2.2 (13,715 genes) and China the Basenji v1.2 (13,712 genes) had more.

Additional kmer analysis of the final assembly [42] yielded 97.32 % (97.2% in chromosomes) and an overall Q-score estimate of 37.5 (38.4 for chromosomes). No sign of retained haplotigs was evident (Supplementary Fig. 3B).

#### Comparison of dingo genomes

We generated a Circos plot [43] to represent the single-nucleotide variants (SNV) and small indel variation between the Alpine and Desert dingo (Fig. 2) using MUMmer4 [44], and sniffles v1.0.11 [45]. In comparison to the autosomes, these plots show low variation on the X chromosome (Fig. 2). To further investigate the low variation, we compared each of the dingoes to CanFam4 (Supplementary Fig. 4, Supplementary Table 1). We then generated a conservative consensus set of structural variants (SV) by merging PacBio, and Nanopore SV calls generated with sniffles [45, 46]. Overall, we found ∼half the number of SV and small variants calls relative to Desert dingo than to CanFam4 (32798 v 62524 and 1729790 v 3839712, respectively).

**Figure 2.**
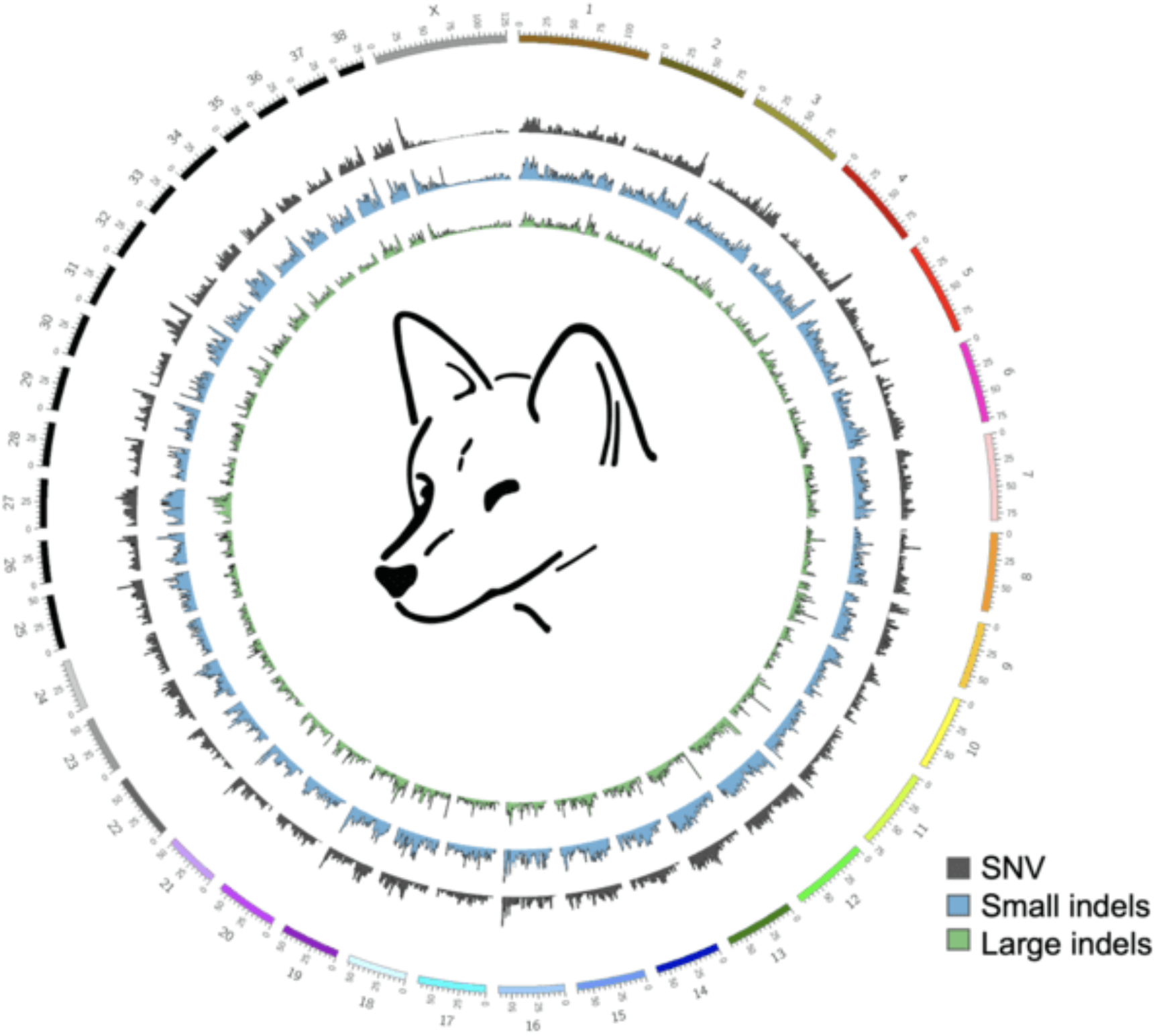
Circos plot comparing Alpine and Desert dingo genomes Plot compares the 38 autosomes and X chromosome of the Alpine and Desert dingo. The plot shows the low variation on the X chromosome compared to the autosomes.

We generated synteny plots using MUMmer plot and GenomeSym [47]. Synteny plots between the dingo genomes show several large-scale chromosomal events. On chromosome 16 there is a 3.45Mb inverted region and a 0.9Mb complex rearrangement (Supplementary Fig. 5). This 3.45Mb inversion does not appear in the wolf or domestic dogs, so we speculate it is unique to the Desert Dingo assembly [6]. The inversion overlaps 60 unique ENSEMBL transcripts and was enriched for gene ontology terms of cellular metabolic processes, including glycolysis and glucose metabolism [6]. Also, on Chromosome 16, the 0.9Mb complex rearrangement occurs between 55 – 57 Mb downstream (Supplementary Fig. 5). Additional structural events include small inversions on Chromosome 11 and on Chromosome 25 (Supplementary Fig. 5). On the X chromosome, there appear to be multiple small nonsyntenic regions (Supplementary Fig. 5); however, further examination of these apparent differences is required to establish whether they are true biological differences or assembly artifacts.

In parallel, we used GeMoMa gene predictions [48] to investigate chromosomal level events. Like the synteny analyses, this approach revealed a large inversion and a disordered region on chromosome 16 as well as smaller inversions on Chromosomes 11 and 25. We also found two structural events on chromosome 26 (Supplementary Fig. 6) containing mostly short genes that are not perfectly conserved (Supplementary Fig. 5F). A MUMmer4 nucmer alignment plot [44] for chromosome 26 corroborated these events (Supplementary Fig. 6)

The Alpine and Desert dingo both have a single copy pancreatic amylase gene (AMY2B) on Chromosome 6. The Alpine dingo assembly does not include a 6.4kb long LINE that was previously reported in the Desert dingo [6].

#### Phylogenetic analyses

All 39 full-length chromosomes in the final assembly were aligned to the corresponding chromosomes in nine published canine *de novo* genome assemblies [6, 27, 28, 36, 38-41]). SNVs and small indels (deletions and insertions <50bp) were called using MUMmer4 call-SNPs module for all possible pairings (Supplementary Table 2). Distance matrices were generated from the inter-canid differences in SNVs and indels and then transformed to WA distance [6, 49]. Fig. 3AC show the phylogenetic tree from SNVs and indels respectively. Both figures show strong support for monophyly of dingoes and dogs relative to the wolf.

**Figure 3.**
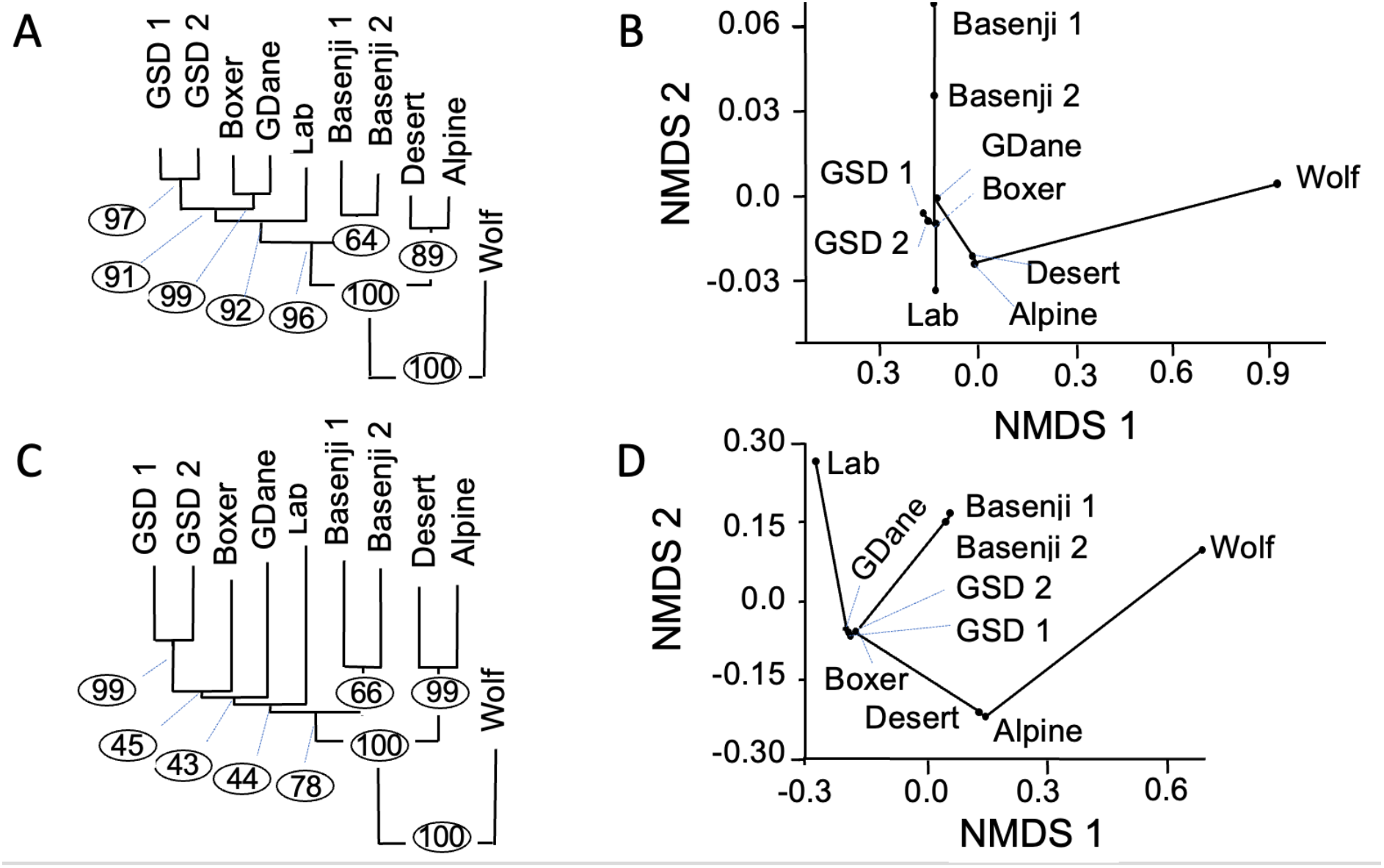
Phylogenetic and ordination analyses of nuclear DNA from SNVs and indels from 10 canines. (A) Phylogenetic tree from SNVs. Branch length proportional to the number of changes and bootstrapping percentage in circles. (B) Ordination analyses from SNVs showing first two axes from non-metric multidimensional scaling (NMDS). (C) Phylogenetic tree from indels. Branch length proportional to the number of changes and bootstrapping percentage in circles. (D) Ordination analyses from indels showing the first two axes from non-metric multidimensional scaling (NMDS). Abbreviations: Lab – Labrador; GSD – German Shepherd Dog; GDane – Great Dane; Wolf — Greenland wolf

These figures also strongly support the hypothesis that dingoes are the sister group to domestic dogs. Fig. 3BD show the ordination analyses from SNVs and indels, respectively. Scores for the taxa calculated from the largest two axes (Axis 1 and Axis 2) describe 75.6% of the variance in SNV’s and 73.2% of the variance in indels (Fig. 3BD).

#### Mitochondrial genome

##### Genome assembly workflow

A 46,192 bp contig from the assembly mapped onto the CanFam reference mtDNA. It constituted a repeat of approximately 2.76 copies of the mtDNA. Following additional polishing and circularization, a final 16,719 bp mtDNA genome was extracted and has been uploaded to GenBank (OP476512).

##### Comparison of dingo mtDNA genomes

When the mtDNA genome of Alpine dingo Cooinda is compared with that of Desert dingo there is a single 10bp SV in the control region that highlights the repeat number difference. In the former, there are 28 repeats (RCGTACACGT) ACGTACGCGCGT, while in the latter, there are 29. Potentially the R(G or A) could represent heteroplasmy [50] that may be further studied with single cell sequencing approaches [51]. Folding this region [52] shows that increasing repeat number increases stem length and overall stability (Supplementary Fig. 7).

Next, we conducted a network analysis in Popart [53] to determine whether the mtDNA of dingo Cooinda fell within the previously described dingo southeastern or northwest clade (Fig. 4) [19, 22]. We included dingo mtDNA from four previous studies, a New Guinea Singing Dog, and an ancient Iron Age dog from Taiwan [6, 22, 54-56]. There were 89 segregating sites and 32 parsimony informative sites in the dataset. Predictably, there were no differences between the mtDNA genome of Cooinda and that previously published from her brother Typia [54]. Further, as expected, Cooinda and Typia mtDNA clustered with samples that had previously been collected from the Alpine region (Fig. 4). Somewhat unexpectedly, the mtDNA from Sandy the dingo found in the desert [6] did not cluster with dingoes from the northwest clade but was closer to canids in the southeastern clade (Fig. 4). This relationship could imply the introgression of Alpine alleles into the Sandy genome however further work would be needed to confirm this.

**Figure 4.**
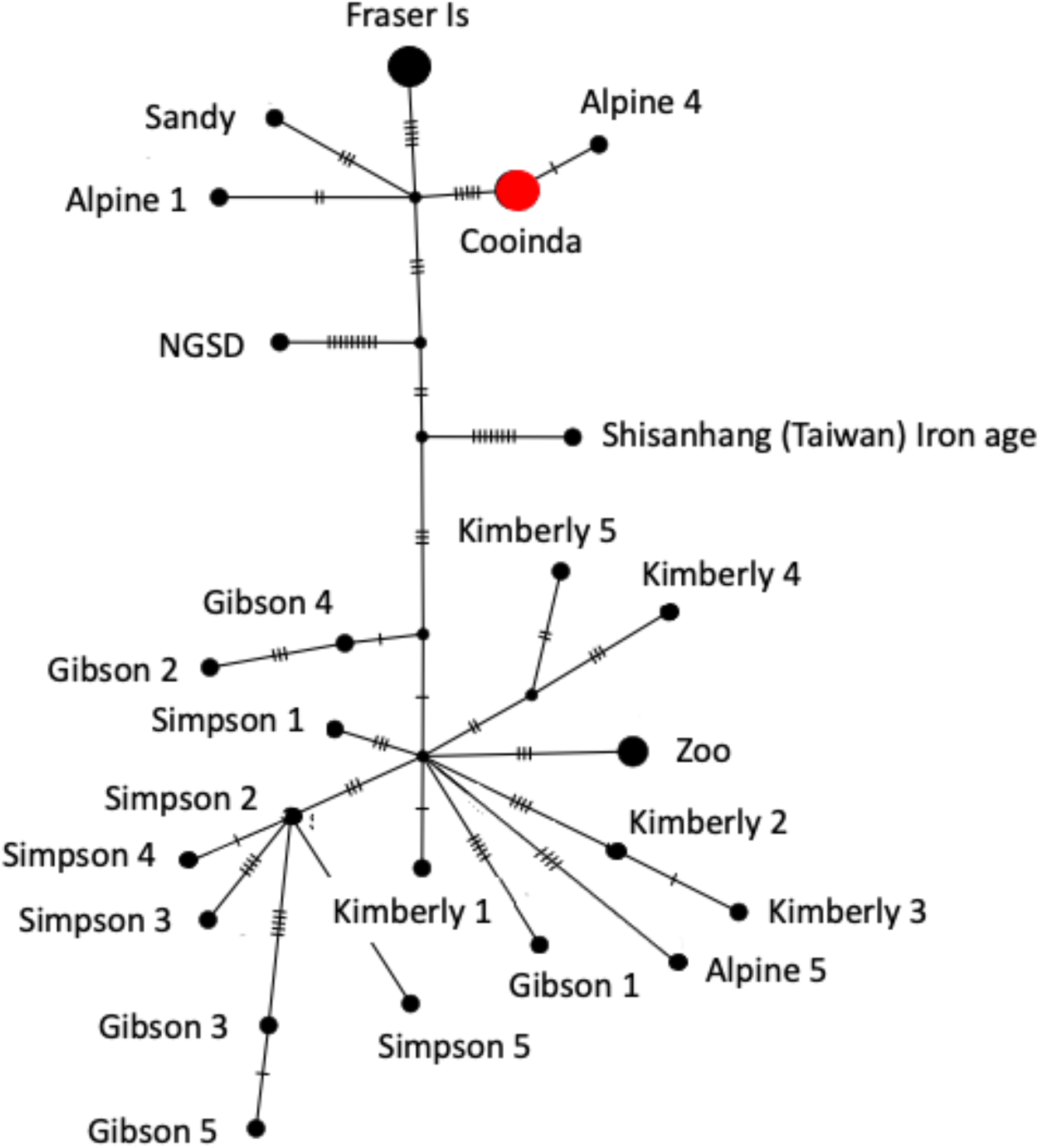
Neighbor-joining network analysis from mtDNA. The size of the circle represents the number of identical sequences and small cross lines the number of SNPs on each branch. The analyses show that dingo Cooinda is in the southeastern clade. Cooinda represents Alpine dingo Cooinda sequenced here, as well as Alpine 2, Alpine 3 [22], MH035670 [55], and Typia [57]. Fraser Is represents the Fraser Island 1-5 samples [22]. Zoo represents three dingoes from the New Zealand Zoo [55]. Shisanhang (Taiwan) is one of two samples from the region and is considered the root of the network [19].

##### DNA methylome

To explore the regulatory landscape of dingo Cooinda, we performed whole genome bisulfite sequencing [58] on genomic DNA extracted from whole blood. In concordance with other adult vertebrates [59, 60], the Cooinda genome displays a typical bimodal DNA methylation pattern. Over 70% of CpG dinucleotides are hypermethylated (levels higher than 80%), and 5% of CpG dinucleotides hypomethylated (methylated at 20% or lower) (Supplementary Fig. 8A).

Next, to determine the number and genomic distribution of putative regulatory regions, we segmented the methylome into unmethylated regions (UMRs) and low-methylated regions (LMRs) using MethylSeekR [61]. UMRs are fully unmethylated and largely coincide with CpG island promoters, whereas LMRs display partial DNA methylation, characteristic of distal regulatory elements such as enhancers in other mammalian models [62]. MethylSeekR analysis identified ∼ 19,000 UMRs and ∼44,000 LMRs in line with previously reported numbers of promoters and enhancers (e.g., human: ∼18,000-20,000 UMRs and 40,000-70,000 LMRs; mouse: ∼17,000-19,000 UMRs and 55,000-90,000 LMRs) [61, 63] (Supplementary Fig. 8BC).

To establish whether proximal gene regulatory regions in the dingo Cooinda genome display different methylation states in the Desert dingo, we converted Cooinda UMR coordinates from Cooinda to the Desert dingo genome assembly using LiftOver (see Methods). Next, we calculated average DNA methylation at Cooinda UMRs and their corresponding lifted-over regions in the Desert dingo genome. We found two UMRs in the Cooinda dingo were hypermethylated in the Desert dingo. These regions overlapped gene bodies of glucagon receptor gene GCGR and histone deacetylase HDAC (Supplementary Fig. 8DE). GCGR is on chromosome 9 and has a single transcript. This transcript is 99.8% identical at the amino acid level between the dingoes. HDAC4 occurs on chromosome 25 and has 12 transcripts with all 12 transcripts being 100% identical at the amino acid level. Further studies are needed to determine the functional significance of the observed differences in DNA methylation.

Altogether, this data provides a genome-wide resource for the putative gene regulatory regions in the Alpine dingo genome, which will be instrumental for future studies.

##### Morphology

###### Skull Morphometrics

Cranial morphology (Supplementary Fig. 9A), quantified using 3D geometric morphometric landmarks, is that of a typical adult female Alpine dingo (Fig. 5). Within the morphospace defined by the principal components explaining the greatest variation between specimens (PC1, PC2), dingo Cooinda’s position is clearly within the Alpine cluster (Fig. 5A). Alpine and Desert dingoes are most clearly differentiated from one another along PC1 (15.70%), for which increasing values describe crania with relatively shorter and broader rostra, shallower orbitals with broader zygomatic arches at the glenoid fossa, prominent and anteriorly-positioned frontals, a higher cranial vault, and prominent sagittal cresting tending to terminate in a high, posteriorly-positioned occiput (inion). Positive values along PC2 (10.60%) mainly denote relatively gracile crania with posteriorly-angled frontals, poorly-developed sagittal cresting, downward-sloping posterior calvarium and a low occipital termination. The sampled Alpine and Desert groups exhibit a near-identical range of PC2 values. As the development of the sagittal cresting, calvarium shape and occipital prominence are related to age and sex, with these traits tending to be more robust and well-developed in males and older dingoes [64], the shared PC2 values across Alpine and Desert groups likely reflect related demographic variation within the respective populations. Within each population (Alpine, Central Desert, Western Desert), males and females overlapped in in their position along PC2 (Supplementary Fig. 9), indicating an absence of strong dimorphism associated with the major axes of shape variance. Despite considerable overlap, PC2 scores tended to be lower in females compared to males in the Alpine and Western Desert populations (see Supplementary Fig. 9, Supplementary Table 3).

**Figure 5.**
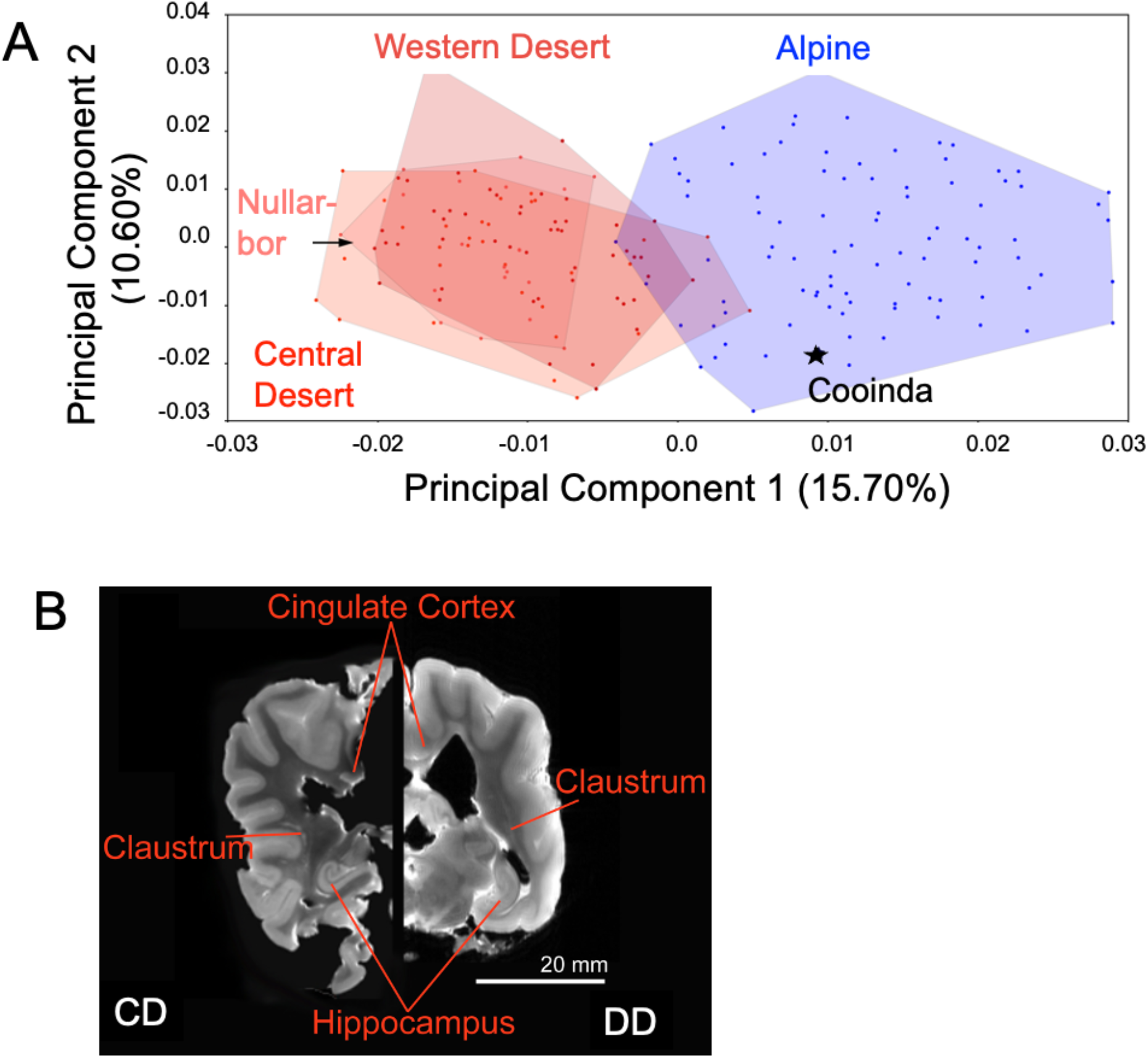
Morphometrics and brain image of Cooinda from the Bargo Dingo Sanctuary, NSW, Australia. (A) Principal Component ordination of geometric morphometric cranial shape data indicating Cooinda’s position in relation to Alpine and Desert dingoes. Blue represents Alpine dingoes, and the red hues indicate dingoes from different Deserts that are broadly overlapping. Dingoes from the Nullarbor overlap most with those from the Alpine region. There is no overlap of dingoes from the Central desert with Alpine dingoes.(B) Brain image, showing a hemispheric comparison of slices generated by Magnetic Resonance (MR) imaging of Cooinda dingo (CD) and a similar-sized domestic dog (DD). Morphometrics and brain image of Cooinda from the Bargo Dingo Sanctuary, NSW, Australia.= (A) Principal Component ordination of geometric morphometric cranial shape data indicating Cooinda’s position in relation to Alpine and Desert dingoes. Blue represents Alpine dingoes, and the red hues indicate dingoes from different Deserts that are broadly overlapping. Dingoes from the Nullarbor overlap most with those from the Alpine region. There is no overlap of dingoes from the Central desert with Alpine dingoes.(B) Brain image, showing a hemispheric comparison of slices generated by Magnetic Resonance (MR) imaging of Cooinda dingo (CD) and a similar-sized domestic dog (DD).

The regression of cranial shape (Procrustes shape variables) on log centroid size (Procrustes shape variables ∼ log(centroid size)) revealed that size contributed significantly to shape variance in the sample (3.91% variance, p <0.001). Size was found to have a non-significant effect on the morphological trajectory described by PC1, which separates Alpine and Desert dingo populations (Fig. 1C), with only 1.23% of related shape-change predicted by centroid size (p = 0.124). Conversely, size predicted 19.88% of shape-change associated with PC2 (p < 0.0001). Alpine and Desert dingo populations share overlapping scores along PC2, and variation along this axis reflects intra-population variability in demographic makeup (age, sex) that should be expected within a natural population. As such, size differences play very little to no role in determining Cooinda’s morphological relationship to Desert dingoes but are important to her position in the Alpine group (Supplementary Fig. 10BC). The low proportion of variation captured in each principal component is a previously-noted feature of the dingo cranial landmark dataset [65] and is unrelated to allometry.

###### Brain imaging

To supplement the morphological data, we quantified brain size. Using a thresholding approach, we used the software 3D Slicer [66] to segment the whole brain as the region of interest. Despite the canids being of very similar size the dingo brain (75.25cm3) was 20% larger than the dog brain (59.53 cm3) (Fig. 5B).

## Discussion

Domestication has received much attention from diverse fields, reflecting the complexity of the process and variation in its duration and intensity [5]. A notable gap in our understanding of the principles of domestication has been the identification of a model system to test Darwin’s two-step predictions [2]. Here we provide the necessary groundwork to explore the potential for dingoes to be a functional and evolutionary intermediate between wild wolves and domestic dogs. One alternate hypothesis is that the process of domestication does not proceed in a stepwise manner [4], but is continual process that represents an intensification of the relationship between a wild species and humans [5].

In this study we compare our high-quality chromosome-level *de novo* assembly of the dingo Cooinda genome with that of the Desert dingo [6], seven domestic dogs [27, 28, 36, 38-40] and the Greenland Wolf [41]. Relative to the wolf and the domestic breeds the Australasian dingo ecotypes are monophyletic. Future studies may include ancient dingo and south east Asian specimens [3], the New Guinea Singing dog [4] and Chinese indigenous dogs [4].

Ancient specimens have potential to give insight into the evolutionary history of dingoes [3] and further instruct the influence of domestic dog admixture [17]. New Guinea Singing Dog may be the sister group to a monophyletic dingo lineage or perhaps more closely related to the Alpine ecotype as suggested by the mtDNA network analyses [19] and cranial shape studies [65]). Inclusion of Chinese indigenous dogs will facilitate determination of the relationships among crown domestic dog breeds [4] and thereby facilitate determination of the divergence date of dingoes and modern dogs.

Multiple large scale chromosomal inversions occur between the two dingo assemblies. There are two large rearrangements on chromosome 16 and likely structural events on Chromosomes 11, 25 and 26 (Supplementary Figs 7, 8). It is also possible that there are multiple small inversions on the X chromosome. It is important to determine the frequency of these events and whether breakpoints affect any regulatory regions or protein coding genes.

Inversions may maintain locally adapted ecotypes, while breakpoints may disrupt regulatory regions or protein coding genes. Hager et al. [67] discovered a 41-megabase chromosomal inversion that characterized defining traits of deer mice (*Peromyscus maniculatus*) and implicated divergent selection in maintaining distinct ecotypes in the wild despite high levels of gene flow. An inversion disrupting FAM134b has been associated with sensory neuropathy in Border Collie dogs [68].

There is a single copy of AMY2B in both dingo genomes; however, they differ by a 6.4 kb retrotransposon insertion present in the Desert dingo. As the retrotransposon is absent in the Greenland wolf and Alpine dingo it would seem likely that the retrotransposon has inserted into the Desert dingo and domestic dog lineages independently. LINE elements can generate duplications through an RNA intermediate and have been associated with amylase expansions in a range of species from humans to mice and rats to dogs [69, 70]. A 1.3kb canid-specific LINE element in domestic dogs is associated with each amylase copy [70].

This expansion is predicted to increase the ability to digest starch [6, 71]. Field et al. [28] compared the influence of *AMY2B* copy number on the microbiomes of dingoes and German Shepherd dogs. They observed distinct and reproducible differences that they hypothesized may influence feeding behaviors. Further studies on *AMY2B* may be fruitful as copy number may be an ecologically relevant mechanism to establish the role of a canid in the ecosystem.

Both dingo ecotypes exhibited low variation on the X chromosome, although it could be argued that variation along the chromosome is not uniform (Fig. 2). Theoretical models predict that genes on the X chromosome can have unusual patterns of evolution due to hemizygosity in males. Sex chromosomes are predicted to exhibit reduced diversity and greater divergence between species and populations compared to autosomes due to differences in the efficacy of selection and drift in these regions [72, 73]. In canids, Plassais et al. [74] show genetic variation in three genes on the X chromosome is strongly associated with body size. Further studies of genetic variation of genes on the X chromosome within and between ecotypes are likely informative.

We integrate the mtDNA genome assembly data with that previously collected from 29 canids in Australasia [6, 22, 54-56]. The mitochondrial genome has been used to infer historical events in various species including canids, but the D-loop region has been difficult to align. Here we show that the region can be folded to increase structural stability with repeat number (Supplementary Fig. 8AB). We found 28, 10-bp repeats in dingo Cooinda compared to 29 in the Desert dingo. The function of the proposed structures is unknown.

Still, folding the region into an extended repeat-dependent stem is expected to decrease the time the DNA in the D-loop is single-stranded during replication. More speculatively, the structure may have a regulatory function that influences mitochondrial bioenergetics and the evolution of mtDNA [75]. Björnerfeldt et al. [76], found that domestic dogs have accumulated nonsynonymous changes in mitochondrial genes at a rate faster than wolves implying a relaxation of selective constraint during domestication.

Phylogenetic and network analyses show that dingo Cooinda has the dingo southeastern Australian mtDNA type of the canine A1b4 subhaplogroup. This southeastern type has been proposed to originate in southern China and includes dogs from Papua New Guinea [19, 22]. Based on mtDNA data, Zhang et al. [19] propose that the TMRCA for most dingoes dates to 6,844 years ago (8,048–5,609 years ago). This estimate is about 3,000 years older than the first known fossil record [77] suggesting that at least two dingo mtDNA haplotypes colonized Australia or older fossil records of dingoes in Australia have yet to be found.

Next, we compare the regulatory landscape of Cooinda dingo with that previously published for the Desert dingo. In comparison to the Alpine dingo, the glucagon receptor gene GCGR and HDAC4 are hypermethylated in the Desert dingo suggesting the potential for dietary or immune differences between ecotypes. Highly methylated gene promoters often indicate a transcriptionally repressed state, while unmethylated gene promoters specify a permissive state [78]. Field et al. [6] previously proposed differences in the feeding behavior of dingoes and wild dogs linked to their *AMY2B* copy number. GCGR is activated by glucagon and initiates a signal transduction pathway that begins with the activation of adenylate cyclase, which in turn produces cyclic AMP. Glucagon is considered the main catabolic hormone of the body and is central to regulating blood glucose and glucose homeostasis [79]. In mice, glucagon has anti-inflammatory properties [80]. HDAC4 is a member of the ubiquitously important family of epigenetic modifier enzymes and has been implicated in processes related to the formation and function of the central nervous system and metabolism. HDAC4 acts as a regulator of pattern-recognition receptor signaling and is involved in regulating innate immune response [81]. In humans, mutations in HDAC4 have been linked with eating disorders [82]. Overlapping conserved Nanopore/PacBio structural variants with these genes identified no variants within GCGR and a single 35bp intronic insertion in HDAC4. The functional impact (if any) of this insertion is unknown.

Dingo Cooinda’s cranial morphology is consistent with the Alpine ecotype from the 20th century. As the first cranial morphological assessment of an Alpine dingo considered to be “pure” by genomic verification, this result is significant in that it suggests that the phenotypic distinctiveness of Alpine dingoes from Desert dingoes is not exclusively the result of recent domestic dog ancestry. Dog admixture has been the predominant explanation given [83] primarily based on the fact that such ancestry is relatively enriched in the southeast region of Australia compared to the north and west [84, 85]. An alternative explanation is that the Alpine and Desert dingoes represent distinct evolutionary lineages. Koungoulos [65] suggested that the cranial shape of Alpine and other southeastern dingoes shares broad similarities with that of New Guinea Singing Dogs and is distinct from the more widespread northwestern lineage [22]. However, these two scenarios are not mutually exclusive. Most introgression likely occurs when a female dingo mates with a male domestic dog. In such cases, extensive backcrossing will not exclude the domestic dog Y. Therefore, examining the Y chromosome of males shown to be pure with the current battery of nuclear-encoded microsatellites will illuminate genetic history. A combination of direct radiocarbon dating, genetic sequencing and morphometric assessment for subfossil material will provide a more confident picture of the nature of change or continuity between ancient and modern Alpine dingoes.

Finally, we supplement our morphological data with magnetic resonance and computed tomography data of Alpine dingo Cooinda’s brain. Her brain was 20% larger than the similarly sized domestic dog, which is consistent with the hypothesis that she was tamed but not domesticated [3] (Fig. 1C). Our brain imaging data are also compatible with prior comparisons that have used endocranial volume as a proxy for brain size, examining a small sample of dingoes (see Geiger et al. [86]) compared to wolves, domestic, basal and archaeological dogs [3]. Endocranial volume in a mixed sample of domestic dogs was shown to be around 30 cm3 smaller than in wolves and jackals [87, 88], which is greater than the 15.7 cm3 difference between the brains of Cooinda and the domestic dog sampled here.

Similarly, brain mass has been shown to be 28.8% smaller in a broad sample (>400) of domestic dogs as compared to wolves [87, 89], which also places the 20% difference between Cooinda and the domestic dog as less pronounced than is seen for comparisons with the wild counterpart (wolf). Brain size reductions are common among domesticated animals compared to their wild counterparts, having been observed across many species, including sheep, pigs, cats, and dogs [87, 90]. Smaller-sized brains, especially size reductions in regions of the forebrain involved in the fight-or-flight response, have been associated with tameness and reductions in fear-based response among domestic animals compared to wild animals [91].

These changes have also been linked to potential reductions in cognitive processing requirements associated with inhabiting anthropogenic environments with lower complexity [92, 93]. Moreover, brain size reductions appear to persist where domestic animals have re-entered a wild environment and exist as feralized animals, at least under certain circumstances [94-96], suggesting that prolonged past exposure to the human niche may be detectable in brain traits. An alternative hypothesis is that differences in brain size is due to environmental adaptation or perhaps Cooinda was an anomaly. Examination of brain size may represent a fruitful pathway for further investigation determining the status of the dingo as a potential feralized animal.

There are at least three possible explanations supporting the existence of two dingo ecotypes (Alpine and Desert). The first is they are ancient Asian lineages that have come into sympatry in Australia. One alternate hypothesis is that a single lineage spread through southeast Asia and then diverged in Australia. There are no major geographical divides in continental Australia, suggesting any differences may reside at the level of biological interactions or they are influenced by climate. In the former case, one possibility is that one or more inversions may maintain the ecotypes [67]. An intriguing alternate hypothesis is that responses to parasites or venomous animals may occur if there are genetic differences in the responses of the ecotypes. In Nigeria, population genomic analyses of 19 indigenous dogs identified 50 positively selected genes including those linked immunity that likely involve adaptations to local conditions [97]. Experimentally it has been shown that adaptation to different parasites or snakes can influence the invasion success of three-spined sticklebacks (*Gasterosteus aculeatus*) and may represent a barrier to gene flow, even between closely related connected populations [98]. In Australia, various parasites and venomous animals have broadly similar distributions to the Alpine ecotype, such as the paralysis tick (*Ixodes holocyclus*) and the red-bellied black snake (*Pseudechis porphyriacus*) [99].

## Conclusions

Here we characterize dingo Cooinda and propose that she be considered the archetype for Australasian dingoes. Characterizing an archetype opens potential for testing Darwin’s [2] two-step model of domestication as an alternative to the hypothesis that domestication represents a continuum [5]. Under the scenario that the dingo has been unconsciously selected, we predict genomic signatures of tameness, as an outcome of unconscious selection [100-102]. Morphologically, we predict lowest shape variation in the rostrum and facial skeleton in the wolf (natural selection), intermediate in the dingo (unconscious selection) and highest in domestic breeds (artificial selection) (i.e., rank order wolf< dingo < modern breeds). Wild populations are more likely to show a narrow range of shape variation about a fitness optimum, whereas changed environmental conditions could support and promote the survival of forms that are farther from the adaptive peak. This is evidenced by earlier research that has shown cranial morphological variation in domestic dogs exceeds that exhibited by the Order Carnivora [26]. In terms of brain size, we predict the magnitude of relative brain size difference will be greater between dingoes and modern breeds than between wolves and dingoes (i.e., rank order wolf> dingo >> modern breeds). Brain size reduction is pronounced in artificial selection and associated with the lack of fear avoidance behavior in domesticates [103]. Dingoes do not show domesticate level reductions in ‘fight or flight’ response [29], and our initial data appear to be at least consistent with this based on the relative brain volume we report.

## Methods

### Sampling: Cooinda the dingo

In selecting an animal for the project, it was considered essential to select an individual that represented the Alpine ecotype, which is found around Sydney, New South Wales (NSW). The individual selected was bred at the Dingo Sanctuary Bargo, NSW, approximately 100km west of Sydney, and has been included in multiple previous studies [6, 29]. Cooinda is the litter sister to Typia from whom short read data had previously been obtained [54]. Cooinda’s parents (Mirri Mirri and Maka), her brothers Typia and Gunya and her were all ginger in color and determined to be pure by microsatellite testing [104]. Mirri Mirri and Maka were independently found in the Alpine region of New South Wales.

An aim of the study is to link genetic and morphological variation, so we provide a brief description of her here. As is typical of Alpine dingoes Cooinda was stocky in appearance with a brad skull and prominent eyes. She was light ginger in color, with dark brown eyes with white paws and chest (Fig. 1AB). Her double coat was not oily like many modern breed dogs and did not have a dog-like odor when wet. She had a pointed muzzle with a broad skull and hooded erect ears. She could turn her neck 180 degrees in any direction. She had lean muscular legs with a long bottle-shaped bushy tail. She weighed 22kg and stood 46cm at the withers. She did not have dewclaws and came into estrus annually. Dingo Cooinda had a loud and clear howl and did not have a modern-dog bark [105]. Cooinda died in 2019 at 10 years of age.

### Chromosome-level genome assembly

#### DNA extraction and sequencing

Genomic DNA for the Pacific Bioscience Single Molecule Real-Time (SMRT) sequencing was prepared from 2 mL of fresh blood using the genomic-tip 100/G kit (Qiagen, Hilden, Germany). This was performed with additional RNase (Astral Scientific, Taren Point, Australia) and proteinase K (NEB, Ipswich, MA, USA) treatment following manufacturer’s instructions. Isolated gDNA was further purified using AMPure XP beads (Beckman Coulter, Brea, CA, USA) to eliminate sequencing inhibitors. DNA purity was calculated using a Nanodrop spectrophotometer (Thermo Fisher Scientific). Molecular integrity was assessed by pulse-field gel-electrophoresis using the PippinPulse (Sage Science) with a 0.75% KBB gel, Invitrogen 1kb Extension DNA ladder and 150 ng of DNA on the 9hr 10-48kb (80V) program. SMRTbell libraries with 20kb insert size were CLR sequenced on Sequel I machines with 2.0 chemistry. Sequencing included 18 SMRT cells with a total polymerase read length 94.25 Gb.

DNA for the Oxford Nanopore (ONT) PromethION sequencing DNA (1 µg) was prepared for ONT sequencing using the 1D genomic DNA ligation kit (SQK-LSK109, ONT) according to the standard protocol. Long fragment buffer was used for the final elution to exclude fragments shorter than 1000 bp. In total, 119 ng of adapted DNA was loaded onto a FLO-PRO002 PromethION flow cell and run on an ONT PromethION sequencing device (PromethION, RRID:SCR_017987) using MinKNOW (18.08.2) with MinKNOW core (v1. 14.2). Base-calling was performed after sequencing with the GPU-enabled guppy basecaller (v3.0.3) using the PromethION high accuracy flip-flop model with config ‘dna_r9.4.1_450bps_hac.cfg’.

For the 10X Genomics Chromium sequencing, DNA was prepared following the protocol described above for SMRT sequencing. A 10X GEM library was barcoded from high-molecular-weight DNA according to the manufacturers recommended protocols. The protocol used was the Chromium Genome Reagent Kits v2 (Document # CG00043 revision B). QC was performed using LabChip GX (PerkinElmer, MA, USA) and Qubit 2.0 Flurometer (Life Technologies, CA, USA). The library was run on a single lane of a v2 patterned flowcell. Sequencing was performed in 150bp paired-end sequencing mode on a single lane on the Illumina HiSeq X Ten platform with a version 2 patterned flowcell.

For the Bionano optical mapping high molecular weight (HMW) DNA was isolated from fresh blood (stored at 4°C) using the Bionano Prep Blood DNA Isolation Protocol following [28]. HMW DNA (∼190 ng/µL) was labelled (BNG, Part #20351) at DLE-1 recognition sites, following the Bionano PrepTM Direct Label and Stain Protocol (BNG, Document #30206 revision C). Labelled DNA was loaded directly onto Bionano Saphyr Chips (BNG, Part #20319), without further fragmentation or amplification, and imaged using a Saphyr instrument to generate single-molecule optical maps. Multiple cycles were performed to reach an average raw genome depth of coverage of 180X.

For the Hi-C sequencing the assembly was scaffolded to chromosome-length by the DNA Zoo following the methodology described here: www.dnazoo.org/methods. Briefly, an *in situ* Hi-C library was prepared [106] from a blood sample of the same female and sequenced to 29X coverage (assuming 2.6 Gb genome size).

#### Workflow

For the initial assembly, The SMRT and ONT reads were corrected and assembled with the Canu assembler (Canu, RRID:SCR_015880; v1.8.0) [31] with the command “canu correctedErrorRate=0.105 corMhapSensitivity=normal corOutCoverage=100 -p Cooinda -d assembly genomesize=2.3g -pacbio-raw Cooinda_SMRT_ONT_combined.fasta. The resulting contigs were polished with two rounds of the Arrow pipeline, each consisting of aligning the raw SMRT reads to the assembly with pbmm2 (https://github.com/PacificBiosciences/pbmm2) and correcting the sequencing errors using gcpp [32].

The Arrow-polished SMRT/ONT assembly was scaffolded using Alpine dingo 10X linked-reads as in ARCS [107]. The 10X data was aligned using the linked-read analysis software provided by 10X Genomics, Long Ranger, v2.1.6 [108]. Misaligned reads and reads not mapping to contig ends were removed, and all possible connections between contigs were computed keeping best reciprocal connections. Finally, contig sequences were joined, spaced by 10kb with stretches of N’s, and if required reverse complemented.

To further improve the assembly, another round of polishing was performed by aligning the Illumina short reads from the 10X Chromium sequencing to the assembly using minimap2 [109] (v2.16) and correcting the sequencing errors using Racon (Racon, RRID:SCR_017642; v1.3.3) [110].

The Hi-C data was processed using Juicer (Juicer, RRID:SCR_017226) [111], and used as input into the 3D-DNA pipeline [112] to produce a candidate chromosome-length genome assembly. We performed additional curation of the scaffolds using Juicebox Assembly Tools [113].

After scaffolding and correction, all raw SMRT and ONT reads were separately aligned to the assembly with Minimap2 (v2.16) (-ax map-pb/map-ont) [109]. The combined alignments were used by PBJelly (pbsuite v.15.8.24) [114] for one round of gap filling.

Following scaffolding, another round of polishing was done to further improve the assembly. Polishing was performed by aligning the Illumina short reads from the Chromium sequencing to the assembly using Long Ranger v2.2.2 and correcting the SNVs and indels using Pilon (Pilon, RRID:SCR_014731) [33].

The Pilon-polished genome underwent a final scaffold clean-up using Diploidocus as described in Edwards et al. [27] to generate a high-quality core assembly, remove low-coverage artefacts and haplotig sequences, and filter any remaining vector/adapter contamination. This reduced the final number of scaffolds to 632 (780 contigs), including the mtDNA.

Assembly completeness was evaluated using BUSCO v5.2.2 [37] short mode against the Carnivora_ob10 data set (n=14,502) implementing BLAST+ v2.11.0 [115], HMMer v3.3 [116], and Metaeuk v20200908 [117]. “Complete” BUSCO genes with available sequences were compiled across Alpine dingo Cooinda and nine canid genomes (Desert dingo [6], two Basenji’s (China and Wags) [27], two German shepherd dogs (Nala and Mischa) [28, 36], Great Dane [38], Labrador [39], Dog10K Boxer [40], and Greenland Wolf [41]) using BUSCOMP v1.0.1. Additional kmer-based assembly completeness and quality evaluations were performed using Merqury v21.3 [42] from the 10x reads.

#### Chromosome mapping and variation

Chromosome mapping was completed in 2019 using the CanFam v3.1 reference genome downloaded from Ensembl (GCF_000002285.3 [118]). Full length chromosomes were renamed with a CANFAMCHR prefix and used for reference mapping. The final Cooinda Alpine dingo genome assembly was mapped onto the CanFam3.1 reference genome using Minimap2 v2.16 [109] (-x asm5 --secondary=no --cs) to generate PAF output. Scaffolds were assigned to CanFam3.1 chromosomes using PAFScaff v0.2.0 [119] based on Minimap2-aligned assembly scaffold coverage against the reference chromosomes. Scaffolds were assigned to the chromosome with highest total coverage. Scaffolds failing to map onto a chromosome were rated as “Unplaced”.

#### Comparison of Alpine and Desert dingo genomes

To investigate the variation between the dingo ecotypes we used Circos [43]. Circos uses a circular ideogram layout to facilitate the display of relationships between the genomes using ribbons, which encode the position and number of SNV’s, small indels and large indels for each of the 38 autosomes and the X chromosome. SNV and indel numbers were calculated using MUMmer4 ‘show-snp’ script following pairwise alignments [44] (v4.0.0 beta 2).

Synteny plot between the Alpine and published Desert dingo assembly [6] was conducted using GenomeSyn [47]. With GenomeSyn the position of the genome is indicated by a black horizontal ruler with tick marks. Syntenic blocks between the genomes are displayed as light grey regions with white illustrating non-syntenic regions. Inversions are represented by red-brown curves.

We used GeMoMa v1.6.2beta [48] to further investigate whole chromosomal events. Here we mapped genes onto the Alpine Dingo assembly following previously described protocols [28]. Subsequently, we checked the synteny of the genes in the reference genome and the target genome using the module GeMoMa module SynthenyChecker. This module uses the GeMoMa annotation with information for reference gene and alternative to determine the best homolog of each transcript. Comparing the order of genes in the reference and the target genome, it allows to determine breakpoints of chromosomal events.

#### Phylogenetic analyses

All 39 full-length chromosomes in the final assembly were aligned to the corresponding chromosomes in nine published canine *de novo* genome assemblies (Desert dingo [6], two basenjis (China and Wags) [27], two German shepherd dogs (Nala and Mischa) [28, 36], Great Dane [38], Labrador [39], Dog10K Boxer [40], and Greenland Wolf [41]) using MUMmer4 [44]. SNVs and small indels (deletions and insertions <50bp) were called using MUMmer4 call-SNPs module for all possible pairings (Supplementary Table 2). Copy number (CNV) and SVs were also called using svmu (v0.2) [120] however these were not included in the phylogeny. SNV’s and indels were analyzed separately. Distance matrices were generated from the inter-canid differences in SNV’s and indels and then transformed to WA distance [49]. Glazko et al. [49] report WA has better phylogenetic properties against normalization of genome sizes than other coefficients.

Phylogenetic analyses using maximum parsimony were generated from the R-package ‘phangorn’ version 2.8.1 [121]. The analyses were run as unrooted networks to test the hypothesis that the wolf was the outgroup. To test the stability of the nodes, a Bayesian bootstrap was applied to the original distance matrix using the program bayesian_bootstrap (github.com/lmc2179/bayesian_bootstrap) and the phylogenetic analysis was re-calculated. This process was iterated 500,000 times. The consensus phylogenetic trees were rooted on the branch leading to wolf, the values indicate the percentage of times that a node occurred. The Y-axis and branch lengths were rescaled to the original number of differences in SNV’s and indels among the taxa. The retention index that measures the fit of the network to the distance matrix exceeded 94% for all 500,000 trees of SNVs and indels.

Non-metric multidimensional scaling (NMDS) was calculated from the distance matrices and scores for the taxa calculated from the largest two axes. Minimum spanning trees were calculated among the scores in NMDS space. NMDS and minimum spanning trees were calculated in Past 4.04 [122].

#### Mitochondrial genome

##### Genome assembly workflow

A 46,192 bp contig from the assembly mapped onto the CanFam reference mtDNA (NC_002008.4), constituting a repeat of approx. 2.76 copies of the mtDNA. The CanFam mtDNA was mapped onto this contig using GABLAM v2.30 [123] and full-length mtDNA copy with highest similarity to CanFam mtDNA was extracted along with 8 kb each side. PacBio reads were mapped onto this mtDNA contig using minimap2 v2.22 [109] and 10x linked reads mapped using BWA v0.7.17 [124] for polishing with HyPo v1.0.3 [125] (32.7 kb assembly size at 673X coverage). The CanFam mtDNA was re-mapped onto the polished assembly using GABLAM v2.30.5 [123] and a 16,719 bp sequence extracted, starting at position 1 of the CanFam sequence. The mtDNA was annotated with the MITOS2 server [126] for submission to NCBI GenBank (accession: OP476512).

##### Comparison of dingo mtDNA genomes

The mtDNA genome of Alpine dingo Cooinda was compared with the Desert dingo [6]. Direct observation of the D-loop region in the two dingoes suggested there was a 10bp repeat and the canids differed in the number of repeats. Imperfect tandem repeats have previously been reported in canids [50]. The D-loop region in Alpine dingo Cooinda was folded using the program mfold [52] to determine ay underlying structures.

To test whether the mtDNA from dingo Cooinda fell within the previously described SE clade we compared the assembly with 33 other canids, including dogs from New Guinea and Taiwan [6, 22, 54, 55]. In this case multiple large gaps were in some of the ancient samples, so the initial assembly was modified based on the predicted secondary structure folding. A inter neighbor-joining network analysis with α = 0.5 was completed in POPART [53]. A limitation of this analyses is that large sections of multiple mtDNA’s were unknown, so it was not possible to distinguish deletions from missing data. Understanding these differences may be biologically important, particularly if the predicted folding of the D-loop region is biologically significant.

### DNA methylome

#### MethylC-seq library preparation

Genomic DNA was extracted from whole blood using DNeasy Blood & Tissue kit (Qiagen, USA). MethylC-seq library preparation was performed as described previously [127].

Briefly, 1 ug of genomic DNA was sonicated to an average size of 300 bp using a Covaris sonicator. Sonicated DNA was then purified, end-repaired and 3’-adenylated followed by the ligation of methylated Illumina TruSeq sequencing adapters. Library amplification was performed with KAPA HiFi HotStart Uracil+ DNA polymerase (Millenium Science Pty Ltd).

#### MethylC-seq data analysis

The methylome library was sequenced on the Illumina HiSeq X platform (150 bp, PE), generating 377M reads. Sequenced reads in fastq format were trimmed using the Trimmomatic software (ILLUMINACLIP:adapter.fa:2:30:10 SLIDINGWINDOW:5:20 LEADING:3 TRAILING:3 MINLEN:50). Trimmed reads were mapped (GCA_012295265.2_UNSW_AlpineDingo_1.0_genomic.fna genome reference, containing the lambda genome as chrLambda) using WALT with the following settings: -m 10 -t 24 -N 10000000 -L 2000. Mapped reads in SAM format were converted to BAM format; BAM files were sorted and indexed using SAMtools. Duplicate reads were removed using Picard Tools v2.3.0. Genotype and methylation bias correction were performed using MethylDackel (MethylDackel extract dingo_lambda.fasta $input_bam -o $output --mergeContext -- minOppositeDepth 5 --maxVariantFrac 0.5 --OT 10,140,10,140 --OB 10,140,10,140). The numbers of methylated and unmethylated calls at each genomic CpG position were determined using MethylDackel (MethylDackel extract dingo_lambda.fasta $input_bam -o output –mergeContext). Segmentation of hypomethylated regions into CpG-rich unmethylated regions (UMRs) and CpG-poor low-methylated regions (LMRs) was performed using MethylSeekR (segmentUMRsLMRs(m=meth, meth.cutoff=0.5, nCpG.cutoff=5, PMDs = NA, num.cores=num.cores, myGenomeSeq=build, seqLengths=seqlengths(build), nCpG.smoothing = 3, minCover = 5).

Cooinda UMR coordinates were converted to the Desert dingo genome assembly using LiftOver following genomewiki.ucsc.edu pipeline (http://genomewiki.ucsc.edu/index.php?title=Minimal_Steps_For_LiftOver). Briefly, the query (Desert dingo) genome build was split into individual scaffolds using *faSplit* (i). The we performed pairwise sequence alignment of query sequences from (i) against the Cooinda genome build using BLAT, Then, coordinates of .psl files were changed to parent coordinate system using *liftUp* and alignments were chained together using axtChain. Chain files were combined and sorted using *chainMergeSort*; alignment nets were made using *chainNet*.

Finally, liftOver chain file was created using *netChainSubset*. Cooinda UMRs in .bed format were lifted over to Desert dingo genome assembly using created liftOver chain file. Average methylation was calculated for Cooinda UMRs and compared to that of corresponding lifted-over regions in the Desert dingo genome. Cooinda UMRs with >50% methylation increase in Desert dingo genome were considered as hypermethylated in the Desert dingo.

### Morphology

#### Skull Morphometrics

To examine cranial morphology, we obtained a 3D model of Cooinda’s cranium using an Artis Pheno Computed Tomography (CT) Scanner. The skull was damaged slightly when the brain was extracted, so the damaged region (dorsal part of the calvarium) was reconstructed using Blender to reassemble the separated fragment following guidelines for digital specimen reconstruction outlined by Lautenschlager [128] (Supplementary Fig. 10A). Geometric morphometric landmarks (n=45) were collected on the 3D cranial model using Stratovan Checkpoint (Stratovan Corporation, Davis, CA version 2018.08.07) and analyzed with MorphoJ [129], following the landmarking protocol used for dingo crania by Koungoulos [65]. This approach uses 45 landmarks along the left side of the cranium, covering all major anatomical features and regions, excepting a few fragile processes which are frequently lost in prepared specimens (Supplementary Fig. 11; Supplementary Table 4). The cranial landmarks collected on the Cooinda cranium were incorporated into an existing data set comprising 91 Alpine dingoes and 101 Desert dingoes [65] and subject to Procrustes superimposition to remove all non-shape differences, due to translation, rotation and scaling [130]. The resultant Procrustes shape variables were ordinated using Principal Component Analysis (PCA) to assess the cranial morphology of Cooinda in relation to other dingoes. To assess the impact of allometry on cranial shape variation in the sample, a regression of Procrustes shape variables against log centroid size was performed using MorphoJ [129].Residuals were extracted from this regression and ordinated using PCA (see Supplementary Material).

#### Brain imaging

Cooinda’s brain and that of a domestic dog (Kelpie) of the same body size were extracted. Brains of these animals, which died within 2 weeks of each other, were fixed in Sigma-Aldrich 10% Neutral Buffered Formalin (NBF) after extraction and were washed with Gd DTPA (gadolinium-diethylenetriamine pentaacetic acid) solution prior to imaging. Brains were scanned using high-resolution magnetic resonance imaging (MRI). A Bruker Biospec 94/20 9.4T high field pre-clinical MRI system was used to acquire MRI data of a fixed dingo and domestic dog brain. The system was equipped with microimaging gradients with a maximum gradient strength of 660mT/m and a 72mm Quadrature volume coil. Images were acquired in transverse and coronal orientation using optimized 2D and 3D Fast Spin Echo (FSE) and Gradient Echo (MGE) methods. Image resolution was 200×200×500 and 300×300 microns isotropic for type 3D and 2D pulse sequences, respectively. To quantify brain size, we used the open-source software 3D Slicer “Segment Statistics” module [66]. The software considers the pixel spacing and slice thickness set to calculate the volume accurately. The threshold was empirically set to the grayscale intensity 1495, where everything below that is background, and ventricles and everything above that is the brain.

## Acknowledgements

Comments from four reviewers improved the manuscript. We would like to thank Luci Ellem, and Dingo Sanctuary Bargo for providing frequent access to Cooinda. Picture of Cooinda was taken by Luci Ellem. Staff at the Vineyard Veterinary Hospital provided constant encouragement. Mike Archer suggested the usage of the term “archetype” and we thank him for valuable taxonomic discussions. Richard Melvin conformed the purity of Cooinda using microsatellites. We thank Shyam Gopalakrishnan and Simon Ho for discussions and Hauke Koch for assistance with translation. SMRT sequencing was conducted at the Ramaciotti Center for Comparative Genomics at University of New South Wales (UNSW). The ONT, 10X Chromium and Bionano genomics data were collected within the Kinghorn Centre for Clinical Genomics at the Garvan Institute of Medical Research, Sydney, Australia and the Hi-C data at Baylor College of Medicine. The high field pre-clinical MRI system was located at the Biological Resources imaging Laboratory at UNSW. Thanks to Jiaming Song for the GenomeSyn analyses, Mihwa Lee for help with DNA folding and Tim Smith for synteny plots. Bootstrapping was on the Wesleyan computing cluster. Thanks go to the facilities of Sydney Imaging at the University of Sydney, and the expertise of Pranish Kolakshyapati in generating the Artis Pheno CT scans of Cooinda’s cranium. Finally, we thank Sandy Ingelby and Harry Parnaby of the Australian Museum for their assistance in facilitating scans of Cooinda’s cranium.

## Availability of supporting data and materials

The chromosomal assembly is available at GCA_012295265.

The mtDNA and has been submitted to NCBI GenBank (accession: OP476512). The methylation data is available at https://www.ncbi.nlm.nih.gov/geo/query/acc.cgi?acc=GSE212509. The 3D Cranial landmark data are available on Figshare at https://figshare.com/articles/dataset/Cooinda_Alpine_Dingo_3D_Cranial_Landmarks/205230

The raw Dicom data for the magnetic resonance imaging (MRI) of the Alpine dingo and domestic dog brain are available on Figshare at https://figshare.com/articles/dataset/Dicom_data_MRI_Alpine_dingo_and_domestic_dog_bra in/20514693.

## Abbreviations

BLAST: Basic Local Alignment Search Tool
BMG: Bionano Genomics
bp: base pairs
BUSCO: Benchmarking Universal Single-Copy Orthologs
CHD: Canine hip dysplasia
d.p: decimal point
CNV: Copy number variant
gDNA: genomic DNA
GSD: German Shepherd Dog
HMM: hidden Markov model
HME: High Molecular Weight
ONT: Oxford Nanopore Technologies
ORF: open reading frame
PacBio: Pacific Biosciences
PCR: polymerase chain reaction
qPCR: quantitative polymerase chain reaction
RNA-seq: RNA sequencing
s.f.: significant figure
SMRT: single-molecule real time
SNV: single-nucleotide variant
SV: structural variant

## Ethics approval and consent to participate

All experimentation was performed under the approval of the University of New South Wales Ethics Committee (ACEC ID: 16/77B).

## Author contributions

JWOB coordinated the project and wrote the initial draft. MAF performed variation analyses. BDR and RJE performed and assisted with the genome assembly, polishing and KAT analysis. LABW and LGK undertook cranial imaging and LGK collected cranial morphometric data. The DNA Zoo initiative, including OD, AO, EA performed and funded the Hi-C experiment. OD and ELA conducted the Hi-C analyses. BC performed the phylogenomic analyses. JK performed the GeMoMa analyses including gene order predictions. OB and KS performed and funded the whole genome bisulphite sequencing and analysis. Eva Chan and Vanessa Hayes collected the Bionano data and performed the analyses. Rob Zammit obtained the initial blood samples and extracted the brain. All authors edited and approved the final manuscript. All authors edited and approved the final manuscript.

## References

1. Darwin C. On the origin of species. London: John Murray; 1858.

2. Darwin C. The variation of animals and plants under domestication. New York: Orange Judd & Co; 1868.

3. Ballard JWO and Wilson LAB. The Australian dingo: untamed or feral? Front Zool. 2019;16:19. doi:10.1186/s12983-019-0300-6.

4. Zhang SJ, Wang GD, Ma P, Zhang LL, Yin TT, Liu YH, et al. Genomic regions under selection in the feralization of the dingoes. Nat Comm. 2020;11:671. doi:10.1038/s41467-020-14515-6.

5. Vigne JD. The origins of animal domestication and husbandry: a major change in the history of humanity and the biosphere. C R Biol. 2011;334 3:171–81. doi:10.1016/j.crvi.2010.12.009.

6. Field MA, Yadav S, Dudchenko O, Esvaran M, Rosen BD, Skvortsova K, et al. The Australian dingo is an early offshoot of modern breed dogs. Sci Adv. 2022;8:eabm5944.

7. White J. Journal of a voyage to New South Wales : with sixty-five plates of non descript animals, birds, lizards, serpents, curious cones of trees and other natural productions. London: Debrett, J.; 1790.

8. Meyer FAA. Systematisch-summarische Uebersicht der neuesten zoologischen Entdeckungen in Neuholland und Afrika: nebst zwey andern zoologischen Abhandlungen. Leipzig: Dykische Buchhandlung; 1793.

9. Crowther MS, Fillios M, Colman N and Letnic M. An updated description of the Australian dingo (Canis dingo Meyer, 1793). J Zool. 2014;293 3:192–203. doi:10.1111/jzo.12134.

10. Smith BP, Cairns KM, Adams JW, Newsome TM, Fillios M, Deaux EC, et al. Taxonomic status of the Australian dingo: the case for Canis dingo Meyer, 1793. Zootaxa. 2019;4564:173–97. doi:10.11646/zootaxa.4564.1.6.

11. Jackson SM, Fleming PJS, Eldridge MDB, Archer M, Ingleby S, Johnson RN, et al. Taxonomy of the dingo: It’s an ancient dog. Aust Zool. 2021;41 3:347–57.

12. Mayr E. Genetics and the origin of species. New York: Columbia University Press; 1942.

13. Jackson SM, Fleming PJS, Eldridge MDB, Ingleby S, Flannery T, Johnson RN, et al. The dogma of dingoes-taxonomic status of the dingo: a reply to Smith et al. Zootaxa. 2019;4564 1.

14. Jackson SM, Groves CP, Fleming PJS, Aplin KP, Eldridge MDB, Gonzalez A, et al. The wayward dog: Is the Australian native dog or dingo a distinct species? Zootaxa. 2017;4317 2:201–24. doi:10.11646/zootaxa.4317.2.1.

15. Corbett LK. The dingo in Australia and Asia. Sydney: University of New South Wales Press; 1995.

16. Corbett L. The conservation status of the dingo Canis lupus dingo in Australia, with particular reference to New South Wales: threats to pure dingoes and potential solutions. In: Dickman CR and Lunney D, editors. A Symposium on the Dingo Sydney: R Zool Soc NSW; 2001.

17. Corbet L. The Australian dingo. In: Merrick JR, Archer M, Hickey GM and Lee SY, editors. Evolution and biogeography of Australian vertebrates. Oatlands, NSW: Australian Scientific Publishing Ltd.; 2006.

18. Jones E. Hybridisation between the dingo, Canis lupus dingo, and the domestic dog, Canis lupus familiaris, in Victoria: a critical review. Aust Mammal. 2009;31:1–7.

19. Zhang M, Sun G, Ren L, Yuan H, Dong G, Zhang L, et al. Ancient DNA evidence from China reveals the expansion of Pacific dogs. Mol Biol Evol. 2020;37:1462–9. doi:10.1093/molbev/msz311.

20. Savolainen P, Leitner T, Wilton AN, Matisoo-Smith E and Lundeberg J. A detailed picture of the origin of the Australian dingo, obtained from the study of mitochondrial DNA. Proc Natl Acad Sci USA. 2004;101 33:12387–90. doi:10.1073/pnas.0401814101.

21. Gonzalez A, Clark G, O’Connor S and Matisoo-Smith L. A 3000 yeAr old dog burial in Timor-Leste. Aust Archaeol. 2013;76:13–9.

22. Cairns KM and Wilton AN. New insights on the history of canids in Oceania based on mitochondrial and nuclear data. Genetica. 2016;144 5:553–65. doi:10.1007/s10709-016-9924-z.

23. Cairns KM, Brown SK, Sacks BN and Ballard JWO. Conservation implications for dingoes from the maternal and paternal genome: multiple populations, dog introgression, and demography. Ecol Evol. 2017;7 22:9787–807. doi:10.1002/ece3.3487.

24. Cairns KM, Shannon LM, Koler-Matznick J, Ballard JWO and Boyko AR. Elucidating biogeographical patterns in Australian native canids using genome wide SNPs. PLoS One. 2018;13 6:e0198754. doi:10.1371/journal.pone.0198754.

25. Freedman AH and Wayne RK. Deciphering the origin of dogs: from fossils to genomes. Annu Rev Anim Biosci. 2017;5:281–307. doi:10.1146/annurev-animal-022114-110937.

26. Drake AG and Klingenberg CP. Large-scale diversification of skull shape in domestic dogs: disparity and modularity. Am Nat. 2010;175 3:289–301. doi:10.1086/650372.

27. Edwards RJ, Field MA, Ferguson JM, Dudchenko O, Keilwagen J, Rosen BD, et al. Chromosome-length genome assembly and structural variations of the primal Basenji dog (Canis lupus familiaris) genome. BMC Genom. 2021;22 1:188. doi:10.1186/s12864-021-07493-6.

28. Field MA, Rosen BD, Dudchenko O, Chan EKF, Minoche AE, Edwards RJ, et al. Canfam_GSD: De novo chromosome-length genome assembly of the German Shepherd Dog (Canis lupus familiaris) using a combination of long reads, optical mapping, and Hi-C. Gigascience. 2020;9 4:giaa027. doi:10.1093/gigascience/giaa027.

29. Ballard JWO, Gardner C, L. Ellem L, Yadav S and R.I. K. Eye-contact and sociability data suggest that Australian dingoes have never been domesticated. Curr Zool. 2021;68 4:423–32.

30. Sluys R. Attaching names to biological species: the use and value of type specimens in systematic zoology and Natural History collections. Biol Theory. 2021;16:49–61.

31. Koren S, Walenz BP, Berlin K, Miller JR, Bergman NH and Phillippy AM. Canu: scalable and accurate long-read assembly via adaptive k-mer weighting and repeat separation. Genome Res. 2017;27 5:722–36. doi:10.1101/gr.215087.116.

32. PacificBiosciences and GenomicConsensus. https://github.com/PacificBiosciences/gcpp.

33. Walker BJ, Abeel T, Shea T, Priest M, Abouelliel A, Sakthikumar S, et al. Pilon: an integrated tool for comprehensive microbial variant detection and genome assembly improvement. PLoS One. 2014;9 11:e112963. doi:10.1371/journal.pone.0112963.

34. Robinson JT, Turner D, Durand NC, Thorvaldsdottir H, Mesirov JP and Aiden EL. Juicebox.js provides a cloud-based visualization system for Hi-C data. Cell Syst. 2018;6 2:256–8 e1. doi:10.1016/j.cels.2018.01.001.

35. DNAZoo: Alpine dingo assembly at DNA Zoo. http://www.dnazoo.org/.

36. Wang C, Wallerman O, Arendt ML, Sundstrom E, Karlsson A, Nordin J, et al. A novel canine reference genome resolves genomic architecture and uncovers transcript complexity. Commun Biol. 2021;4 1:185. doi:10.1038/s42003-021-01698-x.

37. Simao FA, Waterhouse RM, Ioannidis P, Kriventseva EV and Zdobnov EM. BUSCO: assessing genome assembly and annotation completeness with single-copy orthologs. Bioinformatics. 2015;31:3210–2. doi:10.1093/bioinformatics/btv351.

38. Halo JV, Pendleton AL, Shen F, Doucet AJ, Derrien T, Hitte C, et al. Long-read assembly of a Great Dane genome highlights the contribution of GC-rich sequence and mobile elements to canine genomes. Proc Natl Acad Sci USA. 2021;118 11 doi:10.1073/pnas.2016274118.

39. Player RA, Forsyth ER, Verratti KJ, Mohr DW, Scott AF and Bradburne CE. A novel Canis lupus familiaris reference genome improves variant resolution for use in breed-specific GWAS. Life Sci Alliance. 2021;4 4 doi:10.26508/lsa.202000902.

40. Jagannathan V, Hitte C, Kidd JM, Masterson P, Murphy TD, Emery S, et al. Dog10K_Boxer_Tasha_1.0: A Long-Read Assembly of the Dog Reference Genome. Genes. 2021;12 6 doi:10.3390/genes12060847.

41. Sinding MS, Gopalakrishnan S, Raundrup K, Dalen L, Threlfall J, Darwin Tree of Life Barcoding c, et al. The genome sequence of the grey wolf, Canis lupus Linnaeus 1758. Wellcome Open Res. 2021:310. doi:10.12688/wellcomeopenres.17332.1.

42. Rhie A, Walenz BP, Koren S and Phillippy AM. Merqury: reference-free quality, completeness, and phasing assessment for genome assemblies. Genome Biol. 2020;21 1:245. doi:10.1186/s13059-020-02134-9.

43. Krzywinski M, Schein J, Birol I, Connors J, Gascoyne R, Horsman D, et al. Circos: an information aesthetic for comparative genomics. Genome Res. 2009;19:1639–45. doi:10.1101/gr.092759.109.

44. Marcais G, Delcher AL, Phillippy AM, Coston R, Salzberg SL and Zimin A. MUMmer4: A fast and versatile genome alignment system. PLoS Comput Biol. 2018;14 1:e1005944. doi:10.1371/journal.pcbi.1005944.

45. Sedlazeck FJ, Rescheneder P, Smolka M, Fang H, Nattestad M, von Haeseler A, et al. Accurate detection of complex structural variations using single-molecule sequencing. Nat Methods. 2018;15:461–8. doi:10.1038/s41592-018-0001-7.

46. Waardenberg AJ and Field MA. consensusDE: an R package for assessing consensus of multiple RNA-seq algorithms with RUV correction. PeerJ. 2019;7:e8206. doi:10.7717/peerj.8206.

47. Zhou ZW, Yu ZG, Huang XM, Liu JS, Guo YX, Chen LL, et al. GenomeSyn: A bioinformatics tool for visualizing genome synteny and structural variations. J Genet Genom. 2022; doi:10.1016/j.jgg.2022.03.013.

48. Keilwagen J, Hartung F and Grau J. GeMoMa: Homology-Based Gene Prediction Utilizing Intron Position Conservation and RNA-seq Data. Methods Mol Biol. 2019;1962:161–77. doi:10.1007/978-1-4939-9173-0_9.

49. Glazko G, Gordon A and Mushegian A. The choice of optimal distance measure in genome-wide datasets. Bioinformatics. 2005;21 Suppl 3:iii3–11. doi:10.1093/bioinformatics/bti1201.

50. Savolainen P, Arvestad L and Lundeberg J. mtDNA tandem repeats in domestic dogs and wolves: mutation mechanism studied by analysis of the sequence of imperfect repeats. Mol Biol Evol. 2000;17:474–88. doi:10.1093/oxfordjournals.molbev.a026328.

51. Marshall AS and Jones NS. Discovering cellular mitochondrial heteroplasmy heterogeneity with single cell RNA and ATAC sequencing. Biology (Basel). 2021;10 6 doi:10.3390/biology10060503.

52. Zuker M. Mfold web server for nucleic acid folding and hybridization prediction. Nuc Acids Res. 2003;31 13:3406–15. doi:10.1093/nar/gkg595.

53. Leigh JW and Bryant D. Popart: full-feature software for haplotype network construction. Methods Ecol Evol. 2015;6:1110–6.

54. Freedman AH, Gronau I, Schweizer RM, Ortega-Del Vecchyo D, Han E, Silva PM, et al. Genome sequencing highlights the dynamic early history of dogs. PLoS Genet. 2014;10 1:e1004016. doi:10.1371/journal.pgen.1004016.

55. Greig K, Gosling A, Collins CJ, Boocock J, McDonald K, Addison DJ, et al. Complex history of dog (Canis familiaris) origins and translocations in the Pacific revealed by ancient mitogenomes. Sci Rep. 2018;8 1:9130. doi:10.1038/s41598-018-27363-8.

56. Pang JF, Kluetsch C, Zou XJ, Zhang AB, Luo LY, Angleby H, et al. mtDNA data indicate a single origin for dogs south of Yangtze River, less than 16,300 years ago, from numerous wolves. Mol Biol Evol. 2009;26 12:2849–64. doi:10.1093/molbev/msp195.

57. Thalmann O, Shapiro B, Cui P, Schuenemann VJ, Sawyer SK, Greenfield DL, et al. Complete mitochondrial genomes of ancient canids suggest a European origin of domestic dogs. Science. 2013;342:871–4. doi:10.1126/science.1243650.

58. Urich MA, Nery JR, Lister R, Schmitz RJ and Ecker JR. MethylC-seq library preparation for base-resolution whole-genome bisulfite sequencing. Nat Protoc. 2015;10 3:475–83. doi:10.1038/nprot.2014.114.

59. Meissner A, Mikkelsen TS, Gu H, Wernig M, Hanna J, Sivachenko A, et al. Genome-scale DNA methylation maps of pluripotent and differentiated cells. Nature. 2008;454 7205:766–70. doi:10.1038/nature07107.

60. Bogdanovic O, Smits AH, de la Calle Mustienes E, Tena JJ, Ford E, Williams R, et al. Active DNA demethylation at enhancers during the vertebrate phylotypic period. Nat Genet. 2016;48 4:417–26. doi:10.1038/ng.3522.

61. Burger L, Gaidatzis D, Schubeler D and Stadler MB. Identification of active regulatory regions from DNA methylation data. Nucleic Acids Res. 2013;41 16:e155. doi:10.1093/nar/gkt599.

62. Stadler MB, Murr R, Burger L, Ivanek R, Lienert F, Scholer A, et al. DNA-binding factors shape the mouse methylome at distal regulatory regions. Nature. 2011;480 7378:490–5. doi:10.1038/nature10716.

63. Mo A, Mukamel EA, Davis FP, Luo C, Henry GL, Picard S, et al. Epigenomic signatures of neuronal diversity in the mammalian brain. Neuron. 2015;86 6:1369–84. doi:10.1016/j.neuron.2015.05.018.

64. Gollan K. Prehistoric dingo. Australian National University, Canberra, 1982.

65. Koungoulos. K. Old dogs, new tricks: 3D geometric analysis of cranial morphology supports ancient population substructure in the Australian dingo. Zoomorphology. 2020;139:263–75.

66. Fedorov A, Beichel R, Kalpathy-Cramer J, Finet J, Fillion-Robin JC, Pujol S, et al. 3D Slicer as an image computing platform for the quantitative imaging network. Magn Reson Imaging. 2012;30 9:1323–41. doi:10.1016/j.mri.2012.05.001.

67. Hager ER, Harringmeyer OS, Wooldridge TB, Theingi S, Gable JT, McFadden S, et al. A chromosomal inversion contributes to divergence in multiple traits between deer mouse ecotypes. Science. 2022;377 6604:399–405.

68. Forman OP, Hitti RJ, Pettitt L, Jenkins CA, O’Brien DP, Shelton GD, et al. An inversion disrupting FAM134B Is associated with sensory neuropathy in the Border Collie dog breed. G3. 2016;6 9:2687–92. doi:10.1534/g3.116.027896.

69. Tan S, Cardoso-Moreira M, Shi W, Zhang D, Huang J, Mao Y, et al. LTR-mediated retroposition as a mechanism of RNA-based duplication in metazoans. Genome Res. 2016;26:1663–75. doi:10.1101/gr.204925.116.

70. Pajic P, Pavlidis P, Dean K, Neznanova L, Romano RA, Garneau D, et al. Independent amylase gene copy number bursts correlate with dietary preferences in mammals. Elife. 2019;8 doi:10.7554/eLife.44628.

71. Arendt M, Cairns KM, Ballard JWO, Savolainen P and Axelsson E. Diet adaptation in dog reflects spread of prehistoric agriculture. Heredity. 2016;117 5:301–6. doi:10.1038/hdy.2016.48.

72. Vicoso B and Charlesworth B. Evolution on the X chromosome: unusual patterns and processes. Nat Rev Genet. 2006;7 8:645–53. doi:10.1038/nrg1914.

73. Mank JE, Vicoso B, Berlin S and Charlesworth B. Effective population size and the faster-X effect: empirical results and their interpretation. Evolution. 2010;64 3:663–74. doi:10.1111/j.1558-5646.2009.00853.x.

74. Plassais J, Rimbault M, Williams FJ, Davis BW, Schoenebeck JJ and Ostrander EA. Analysis of large versus small dogs reveals three genes on the canine X chromosome associated with body weight, muscling and back fat thickness. PLoS Genet. 2017;13 3:e1006661. doi:10.1371/journal.pgen.1006661.

75. Basu U, Bostwick AM, Das K, Dittenhafer-Reed KE and Patel SS. Structure, mechanism, and regulation of mitochondrial DNA transcription initiation. J Biol Chem. 2020;295 52:18406–25. doi:10.1074/jbc.REV120.011202.

76. Bjornerfeldt S, Webster MT and Vila C. Relaxation of selective constraint on dog mitochondrial DNA following domestication. Genome Res. 2006;16 8:990–4. doi:10.1101/gr.5117706.

77. Milham PT, P. Relative antiquity of human occupation and extinct fauna at Madura Cave, Southeastern Western Australia. Mankind. 1976;10:175–80.

78. Schubeler D. Function and information content of DNA methylation. Nature. 2015;517 7534:321–6. doi:10.1038/nature14192.

79. Wewer Albrechtsen NJ, Kuhre RE, Pedersen J, Knop FK and Holst JJ. The biology of glucagon and the consequences of hyperglucagonemia. Biomark Med. 2016;10 11:1141–51. doi:10.2217/bmm-2016-0090.

80. Insuela DBR, Azevedo CT, Coutinho DS, Magalhaes NS, Ferrero MR, Ferreira TPT, et al. Glucagon reduces airway hyperreactivity, inflammation, and remodeling induced by ovalbumin. Sci Rep. 2019;9 1:6478. doi:10.1038/s41598-019-42981-6.

81. Yang Q, Tang J, Pei R, Gao X, Guo J, Xu C, et al. Host HDAC4 regulates the antiviral response by inhibiting the phosphorylation of IRF3. J Mol Cell Biol. 2019;11:158–69. doi:10.1093/jmcb/mjy035.

82. Cui H, Moore J, Ashimi SS, Mason BL, Drawbridge JN, Han S, et al. Eating disorder predisposition is associated with ESRRA and HDAC4 mutations. J Clin Invest. 2013;123 11:4706–13. doi:10.1172/JCI71400.

83. Radford CG, Letnic M, Fillios M and Crowther MS. An assessment of the taxonomic status of wild canids in south-eastern New South Wales: phenotypic variation in dingoes. Aust J Zool. 2012;60:73–80.

84. Stephens D, Wilton AN, Fleming PJ and Berry O. Death by sex in an Australian icon: a continent-wide survey reveals extensive hybridization between dingoes and domestic dogs. Mol Ecol. 2015;24 22:5643–56. doi:10.1111/mec.13416.

85. Cairns KM, Crother MS, Nesbit B and Letnik M. The myth of wild dogs in Australia: are there any out there? Aust Mamm. 2020;44:67–75.

86. Geiger M, Evin A, Sanchez-Villagra MR, Gascho D, Mainini C and Zollikofer CPE. Neomorphosis and heterochrony of skull shape in dog domestication. Sci Rep. 2017;7 1:13443. doi:10.1038/s41598-017-12582-2.

87. Balcarcel AM, Geiger M, Clauss M and Sanchez-Villagra MR. The mammalian brain under domestication: discovering patterns after a century of old and new analyses. J Exp Zool B Mol Dev Evol. 2022;338 8:460–83. doi:10.1002/jez.b.23105.

88. Klatt B. Über die veränderung der schädelkapazität in der somestikation. Sitzungsbericht der Gesellschaft naturforschender Freunde. 1912:3.

89. Röhrs M and Ebinger P. Die Berteilung von Hirngrossenunterschieden. Journal of Zoological Systematics and Evolutionary Research. 1978;16:1–14.

90. Kruska D. Mammalian domestication and its effect on brain structure and behavior. In: Jerison H J, and Jerison I, editors. Intelligence and Evolutionary Biology. New York: Academic Press; 1988.

91. Brusini I, Carneiro M, Wang C, Rubin CJ, Ring H, Afonso S, et al. Changes in brain architecture are consistent with altered fear processing in domestic rabbits. Proc Natl Acad Sci USA. 2018;115 28:7380–5. doi:10.1073/pnas.1801024115.

92. Kruska DC. On the evolutionary significance of encephalization in some eutherian mammals: effects of adaptive radiation, domestication, and feralization. Brain Behav Evol. 2005;65 2:73–108. doi:10.1159/000082979.

93. Barrickman NL, Bastian ML, Isler K and van Schaik CP. Life history costs and benefits of encephalization: a comparative test using data from long-term studies of primates in the wild. J Hum Evol. 2008;54 5:568–90. doi:10.1016/j.jhevol.2007.08.012.

94. Rohrs M and Ebinger P. Wild is not really wild: brain weight of wild domestic mammals. Berl Munch Tierarztl Wochenschr. 1999;112 6-7:234–8.

95. Kruska D and M. R. Comparative-quantitative investigations on brains of feral pigs from the Galapagos Islands and of European domestic pigs. Z Anat Entwicklungsgesch. 1974;144:61–73.

96. Lord KA, Larson G and Karlsson EK. Brain size does not rescue domestication syndrome. Trends Ecol Evol. 2020;35 12:1061–2. doi:10.1016/j.tree.2020.10.004.

97. Liu YH, Wang L, Xu T, Guo X, Li Y, Yin TT, et al. Whole-genome sequencing of African dogs provides Insights into adaptations against tropical parasites. Mol Biol Evol. 2018;35 2:287–98. doi:10.1093/molbev/msx258.

98. Erin NI, Benesh DP, Henrich T, Samonte IE, Jakobsen PJ and Kalbe M. Examining the role of parasites in limiting unidirectional gene flow between lake and river sticklebacks. J Anim Ecol. 2019;88 12:1986–97. doi:10.1111/1365-2656.13080.

99. Bradley C. Venomous bites and stings in Australia to 2005. In: Welfare AIoHa, (ed.). Canberra: Australian Government, 2014, p. 119.

100. Gulevich RG and et al. Effect of selection for behavior on pituitary-adrenal axis and proopiomelanocortin gene expression in silver foxes (Vulpes vulpes). Physiol Behav. 2004;82 2-3:513–8. doi:10.1016/j.physbeh.2004.04.062.

101. Heyne HO, Lautenschläger S, Nelson R, Besnier F, Rotival M, Cagan A, et al. Genetic influences on brain gene expression in rats selected for tameness and aggression. Genetics. 2014;198 3:1277–90. doi:10.1534/genetics.114.168948.

102. Matsumoto Y, Nagayama. H., Nakaoka H, Toyoda A, Goto T and Koide T. Combined change of behavioral traits for domestication and gene-networks in mice selectively bred for active tameness. Genes Brain Behav. 2021;20:e12721. doi:10.1111/gbb.12721.

103. Albert FW and et al. A comparison of brain gene expression levels in domesticated and wild animals. PLoS Genet. 2012;8 9:e1002962. doi:10.1371/journal.pgen.1002962.

104. Wilton AN. DNA methods of assessing dingo purity.. Sydney: R. Zool. Soc. N.S.W.; 2001.

105. Deaux EC, Allen AP, Clarke JA and Charrier I. Concatenation of ‘alert’ and ‘identity’ segments in dingoes’ alarm calls. Sci Rep. 2016;6:30556. doi:10.1038/srep30556.

106. Rao SS, Huntley MH, Durand NC, Stamenova EK, Bochkov ID, Robinson JT, et al. A 3D map of the human genome at kilobase resolution reveals principles of chromatin looping. Cell. 2014;159 7:1665–80. doi:10.1016/j.cell.2014.11.021.

107. Yeo S, Coombe L, Warren RL, Chu J and Birol I. ARCS: scaffolding genome drafts with linked reads. Bioinformatics. 2018;34:725–31. doi:10.1093/bioinformatics/btx675.

108. Chromium X: 10X Genomics linked-read alignment,variant calling, phasing, and structural variant calling https://support.10xgenomics.com/genome-exome/software/pipelines/latest/what-is-long-ranger (2020). Accessed 2020.

109. Li H. Minimap2: pairwise alignment for nucleotide sequences. Bioinformatics. 2018;34 18:3094–100. doi:10.1093/bioinformatics/bty191.

110. Vaser R, Sovic I, Nagarajan N and Sikic M. Fast and accurate de novo genome assembly from long uncorrected reads. Genome Res. 2017;27 5:737–46. doi:10.1101/gr.214270.116.

111. Durand NC, Robinson JT, Shamim MS, Machol I, Mesirov JP, Lander ES, et al. Juicebox provides a visualization system for Hi-C contact maps with unlimited zoom. Cell Syst. 2016;3 1:99–101. doi:10.1016/j.cels.2015.07.012.

112. Dudchenko O, Batra SS, Omer AD, Nyquist SK, Hoeger M, Durand NC, et al. De novo assembly of the Aedes aegypti genome using Hi-C yields chromosome-length scaffolds. Science. 2017;356 6333:92–5. doi:10.1126/science.aal3327.

113. Dudchenko O, Shamim MS, Batra SS, Durand NC, Musial NT, Mostofa R, et al. The Juicebox Assembly Tools module facilitates de novo assembly of mammalian genomes with chromosome-length scaffolds for under $1000. bioRxiv. 2018:254797. doi:10.1101/254797.

114. English AC, Richards S, Han Y, Wang M, Vee V, Qu J, et al. Mind the gap: upgrading genomes with Pacific Biosciences RS long-read sequencing technology. PLoS One. 2012;7 11:e47768. doi:10.1371/journal.pone.0047768.

115. Altschul SF, Gish W, Miller W, Myers EW and Lipman DJ. Basic local alignment search tool. J Mol Biol. 1990;215 3:403–10. doi:10.1016/S0022-2836(05)80360-2.

116. Finn RD, Clements J and Eddy SR. HMMER web server: interactive sequence similarity searching. Nucleic Acids Res. 2011;39 Web Server issue:W29–37. doi:10.1093/nar/gkr367.

117. Levy KE, Mirdita M and Soding J. MetaEuk-sensitive, high-throughput gene discovery, and annotation for large-scale eukaryotic metagenomics. Microbiome. 2020;8 1:48. doi:10.1186/s40168-020-00808-x.

118. Hoeppner MP, Lundquist A, Pirun M, Meadows JR, Zamani N, Johnson J, et al. An improved canine genome and a comprehensive catalogue of coding genes and non-coding transcripts. PLoS One. 2014;9 3:e91172. doi:10.1371/journal.pone.0091172.

119. Edwards R: PAFScaff biotools. https://bio.tools/PAFScaff_Pairwise_mApping_Format_reference-based_scaffold_anchoring_and_super-scaffolding. (2020). Accessed Nov 1, 2019.

120. Chakraborty M, Emerson JJ, Macdonald SJ and Long AD. Structural variants exhibit widespread allelic heterogeneity and shape variation in complex traits. Nat Commun. 2019;10 1:4872. doi:10.1038/s41467-019-12884-1.

121. Schliep K, Potts AJ, Morrison DA and Grimm GW. Intertwining phylogenetic trees and networks. Methods Ecol Evol. 2017;8 10:1212–20.

122. Hammer O, Harper DAT and PD. R. PAST: Paleontological software package for education and data ananlysis. Palaeontol Electron. 2001;4:9pp.

123. Davey NE, Shields DC and Edwards RJ. SLiMDisc: short, linear motif discovery, correcting for common evolutionary descent. Nuc Acids Res. 2006;34 12:3546–54. doi:10.1093/nar/gkl486.

124. Li H and Durbin R. Fast and accurate short read alignment with Burrows-Wheeler transform. Bioinformatics. 2009;25 14:1754–60. doi:10.1093/bioinformatics/btp324.

125. Kundu R, Casey J and Sung W-K. HyPo: Super fast & accurate polisher for long read genome assemblies. bioRxiv. 2019:doi: 10.1101/2019.12.19.882506. doi:10.1101/2019.12.19.882506.

126. Donath A, Juhling F, Al-Arab M, Bernhart SH, Reinhardt F, Stadler PF, et al. Improved annotation of protein-coding genes boundaries in metazoan mitochondrial genomes. Nucleic Acids Res. 2019;47 20:10543–52. doi:10.1093/nar/gkz833.

127. Urich MA, Nery JR, Lister R, Schmitz RJ and Ecker JR. MethylC-seq library preparation for base-resolution whole-genome bisulfite sequencing. Nat Protoc. 2015;10 3:475–83. doi:10.1038/nprot.2014.114.

128. Lautenschlager S. Reconstructing the past: methods and techniques for the digital restoration of fossils. R Soc Open Sci. 2016;3 10:160342. doi:10.1098/rsos.160342.

129. Klingenberg CP. MorphoJ: an integrated software package for geometric morphometrics. Mol Ecol Resour. 2011;11 2:353–7. doi:10.1111/j.1755-0998.2010.02924.x.

130. Rohlf F and Slice D. Extensions of the procrustes method for the optimal superimposition of landmarks. Syst Zool. 1990;39.

