## Supplemental Figures and Tables for "The Australasian dingo archetype: *De novo* chromosome-length genome assembly, DNA methylome, and cranial morphology"

### Additional Files

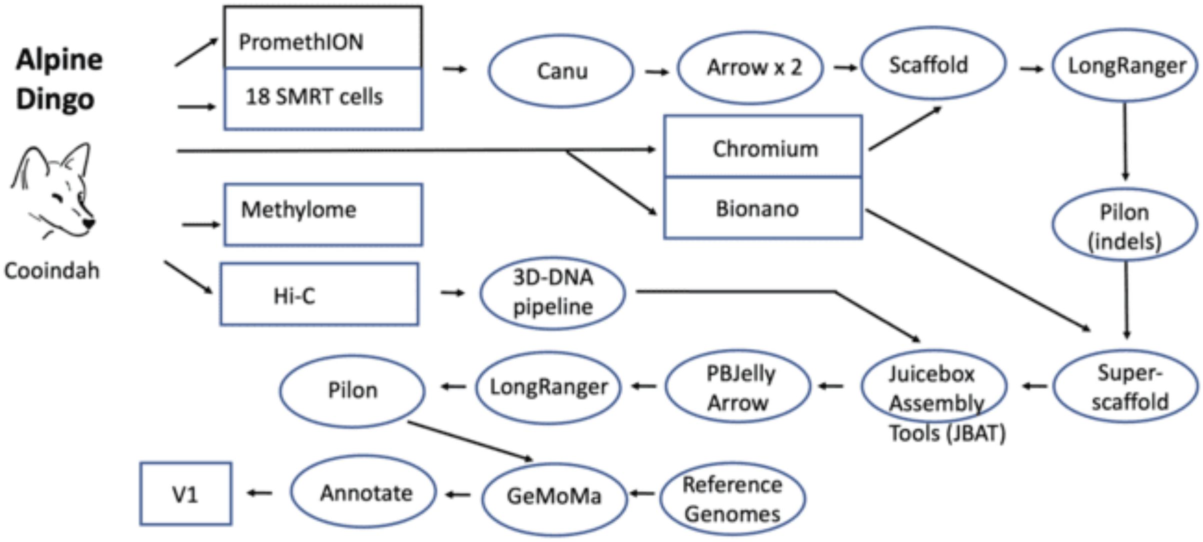


**Supplementary Figure 1 Title.** Schematic overview of project workflow

**Supplementary Figure 1 Legend.** Alpine Dingo Cooinda DNA was derived from blood of a single female from the Dingo Sanctuary Bargo. Sequences were generated on the Pacific Biosciences Sequel instrument (V2 chemistry) and Oxford Nanopore PromethION instrument (guppy bascaller Version 3.0.6+9999d81) to ~30x genome coverage, each, based on a genome size estimate of 2.4 Gb (this estimate is used for all coverage estimates). All long read sequences were assembled with the Canu v1.8 algorithm then error corrected twice using the Arrow genomic consensus polishing module. The assembly was scaffolded with Chromium 10x linked-reads (~41x coverage excluding the barcode) using Long Ranger v2.1.6 using DNA from the same animal. Polishing of the assembly for residual indels was done by aligning the Illumina data with Minimap2 and the Racon algorithm. Single molecule Bionano data (~57x effective coverage) was then used to superscaffold the sequence assembly using DNA extracted from the same canid. For this, single molecule optical maps were first de novo assembled into consensus maps, which were than aligned to the sequence assembly in silico digested with the same labelling enzyme for hybrid scaffolding, using Bionano Solve (v3.2.2_08022018) with RefAligner (7782.7865rel). This assembly was further scaffolded to chromosome-length by DNA Zoo (www.dnazoo.org/methods). Briefly, an *in situ* Hi-C library was prepared from the same individual and sequenced to 29x coverage. The Hi-C data was processed using Juicer [111], and used as input into the 3D-DNA pipeline [112] to produce a candidate chromosome-length genome assembly. We performed additional finishing on the scaffolds using Juicebox Assembly Tools [113]. The assembly was then long-read gap filled with the PBJelly algorithm, and the additional data error corrected using Arrow [32]. The Chromium data was mapped onto the assembly with the Long Ranger v2.1.6 program and the final assembly was then polished using the Pilon algorithm. Of the 2.4 Gb assembled genome, the total assembly N50 contig and scaffold lengths are 23.1 Mb and 64.8 Mb, respectively. The assembled contigs were then aligned to CanFam3.1 for chromosome assignments. Regulatory landscape was characterised by whole genome bisulphite sequencing.

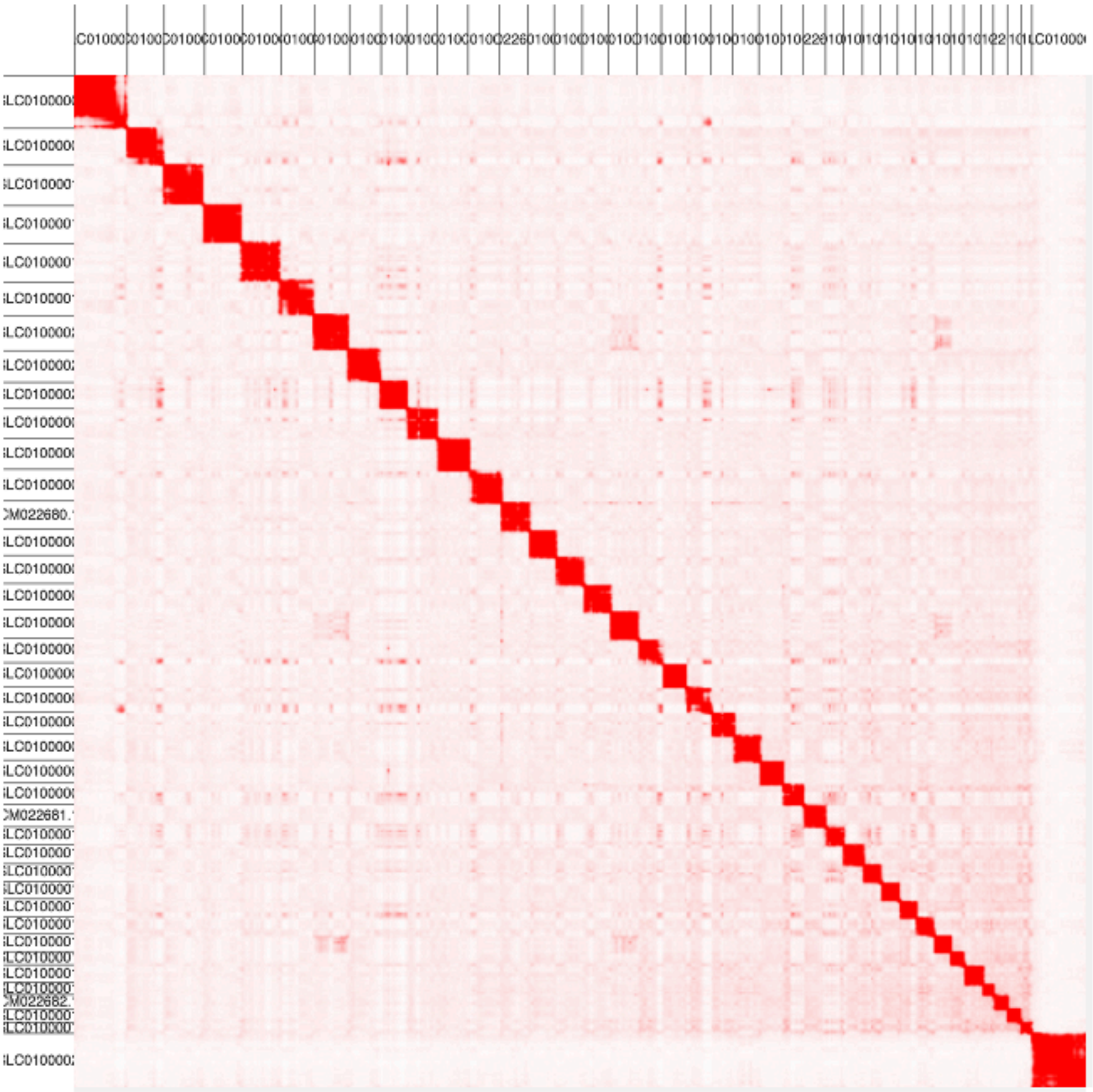

**Supplementary Figure 2 Title:** Alpine dingo assembly after Hi-C correction

**Supplementary Figure 2 Legend:** Contact matrices (visualised in Juicebox.js) after the chromosome-length Hi-C upgrade. The chromosome-length contact map can be viewed at multiple resolutions using Juicebox.js [34] following the link https://tinyurl.com/ycbkezf4.

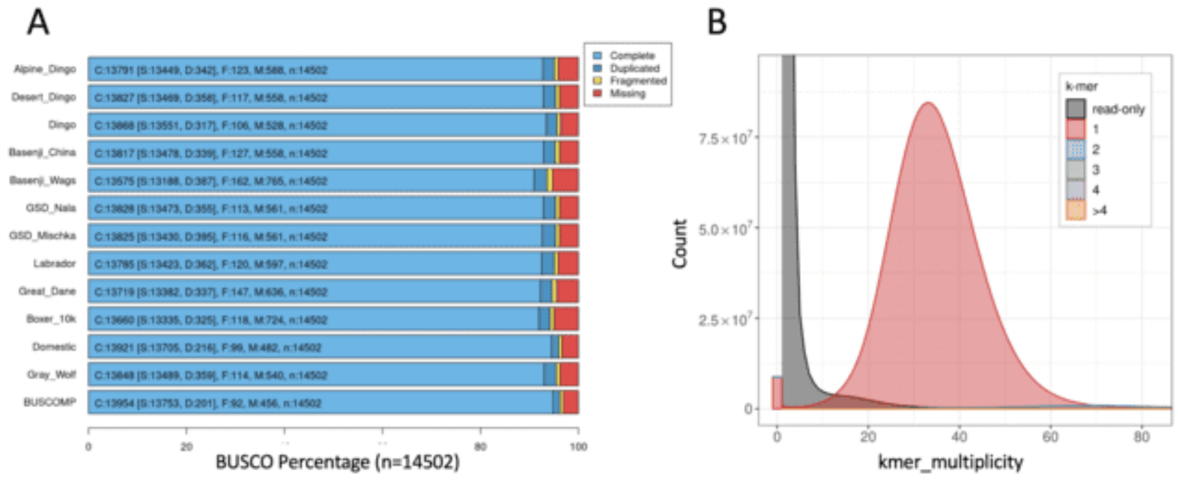


**Supplementary Figure 3 Title:** Assembly statistics

**Supplementary Figure 3 Legend:** (A) BUSCO ratings for Cooinda assembly, compared to CanFam4. Purple, original assembly; Black, scaffolding/polishing steps; Blue, final assembly; Red, CanFam4. Dashed red lines mark CanFam4 statistics.

(B) 10x read kmers frequency distributions for kmers with different assembly copy numbers derived from A Read 1 (16bp barcodes trimmed) and B Read 2 (barcodes not trimmed).

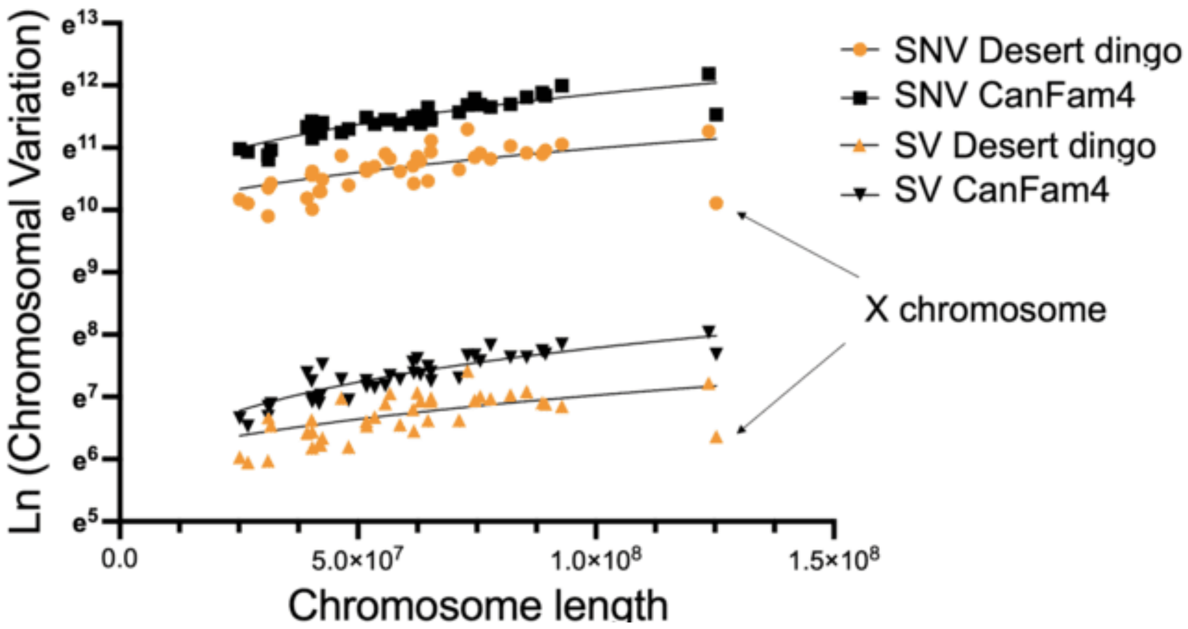


**Supplementary Figure 4:** Genome shows a deficiency of variation on the X chromosome

**Supplementary Figure 4 Legend:** SNV (single nucleotide variant) and SV (structural variation) comparisons show a relative deficiency of variation on the X chromosome. Line represents a regression through the non-transformed data and each point represents one chromosome with the length of the Alpine dingo and SNV's or SV relative to the Desert dingo genome or CanFam4. Y=3.8e-4x+21305, 1.1e-4+31753, 7.2e-5+406.7, 2.5e-5+363.1 with an $r^2$ of 0.37, 0.74, 0.33, 0.77 for SNV Desert dingo, SNV CanFam, SV Desert dingo and SV CanFam, respectively. If the SNV and SV Desert dingo X chromosome data are excluded the $r^2$ of these regressions increases to 0.67 and 0.54, respectively.

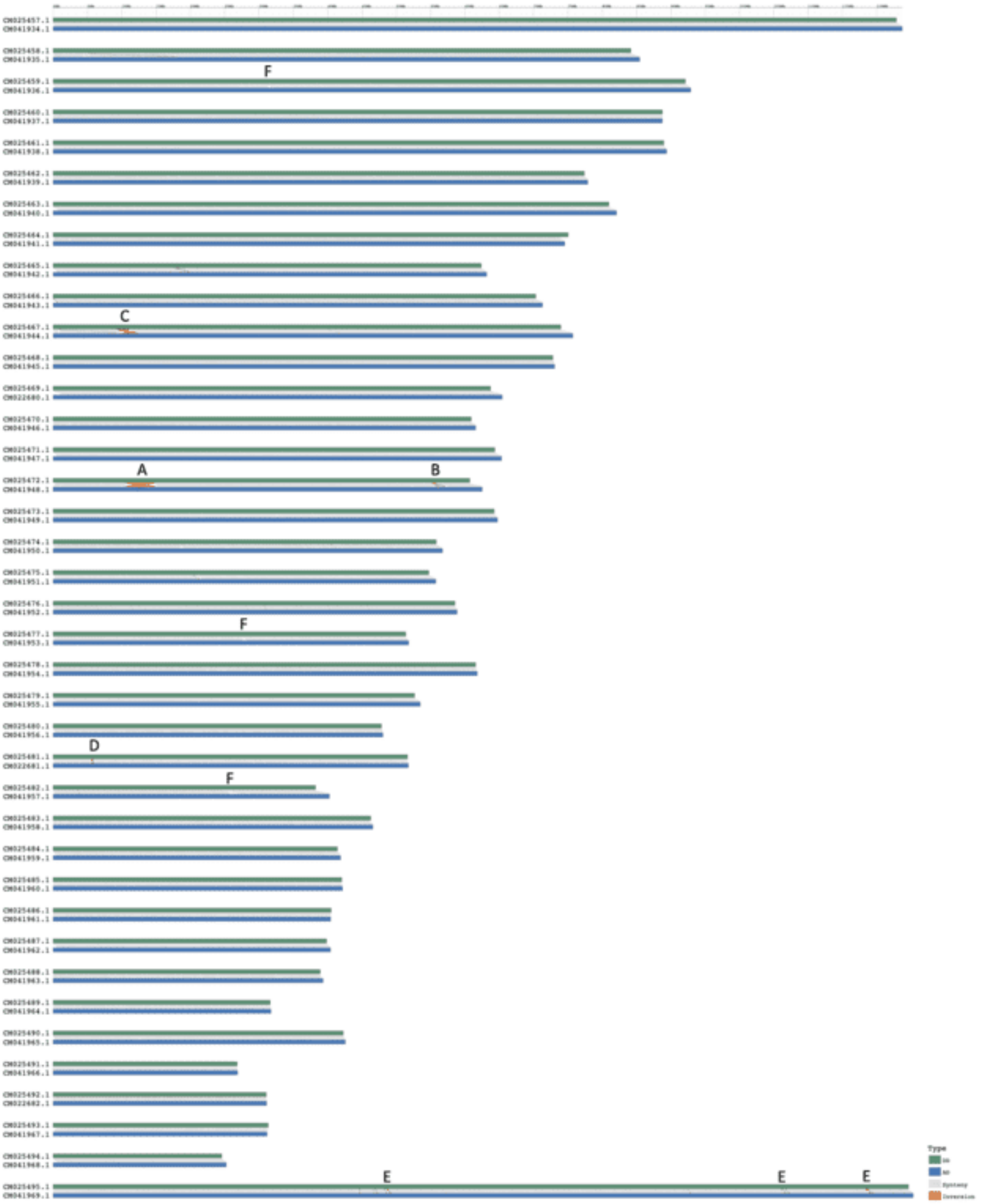


**Supplementary Figure 5:** Synteny analyses

**Supplementary Figure 5 Legend:** Synteny plot of Alpine dingo Cooinda (AD) in blue against Desert dingo Sandy in orange (DD). A. Shows the 3.45Mb rearrangement on Chromosome 16. B. Shows the complex rearrangement between 55-57 Mb downstream on Chromosome 16. C Smaller inversion on Chromosome 11. D. Small inversion on

Chromosome 25. E. Multiple possible small inversions on X chromosome. Other smaller rearrangements are possible. F. Possible duplication like events.

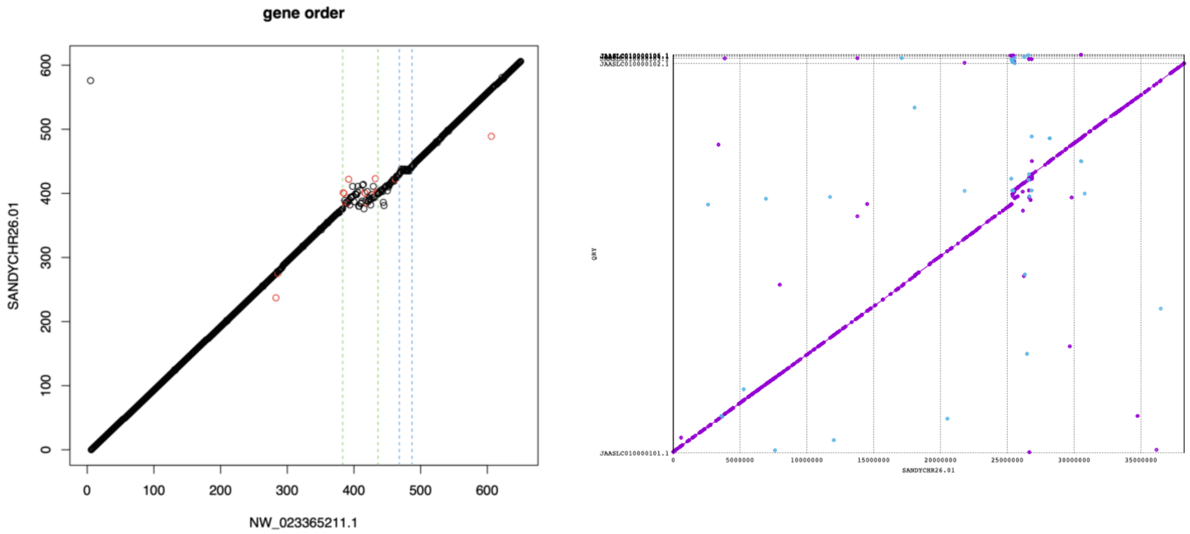


**Supplementary Figure 6:** Gene order plot comparing Chromosome 26 for Cooinda the Alpine Dingo (X-axis) and Desert Dingo Sandy (Y-axis) using GenomeSym (left) and MUMmer (right).

**Supplementary Figure 6 Legend:** In GeMoMa plot (left) the green and the blue dashed lines indicate the two structural events on chromosome 26 of Cooinda. The same region is shown using MUMmer (right).

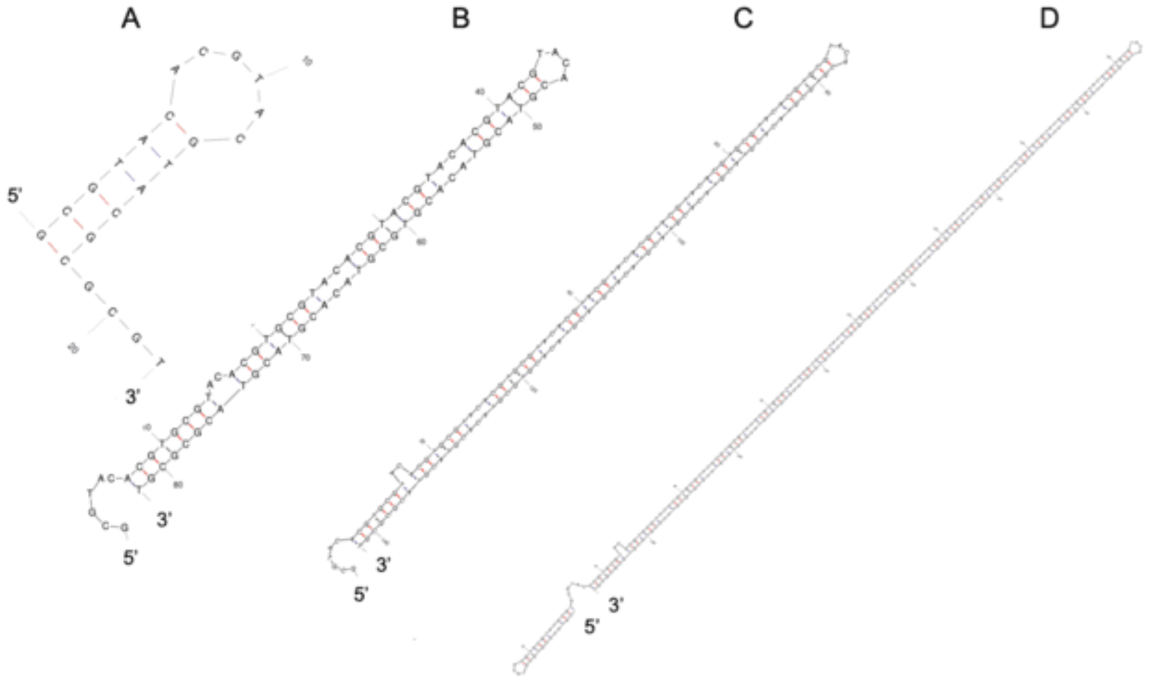


**Supplementary Figure 7:** Possible folding of 10bp repeats in D-loop region

**S Supplementary Figure 7 Legend:** (A) 1 repeat, ΔG=-4.68, (B) 7 repeats ΔG=-29.07, (C) 13 repeats ΔG=-48.21, (D) 28 repeats ΔG=-97.71.

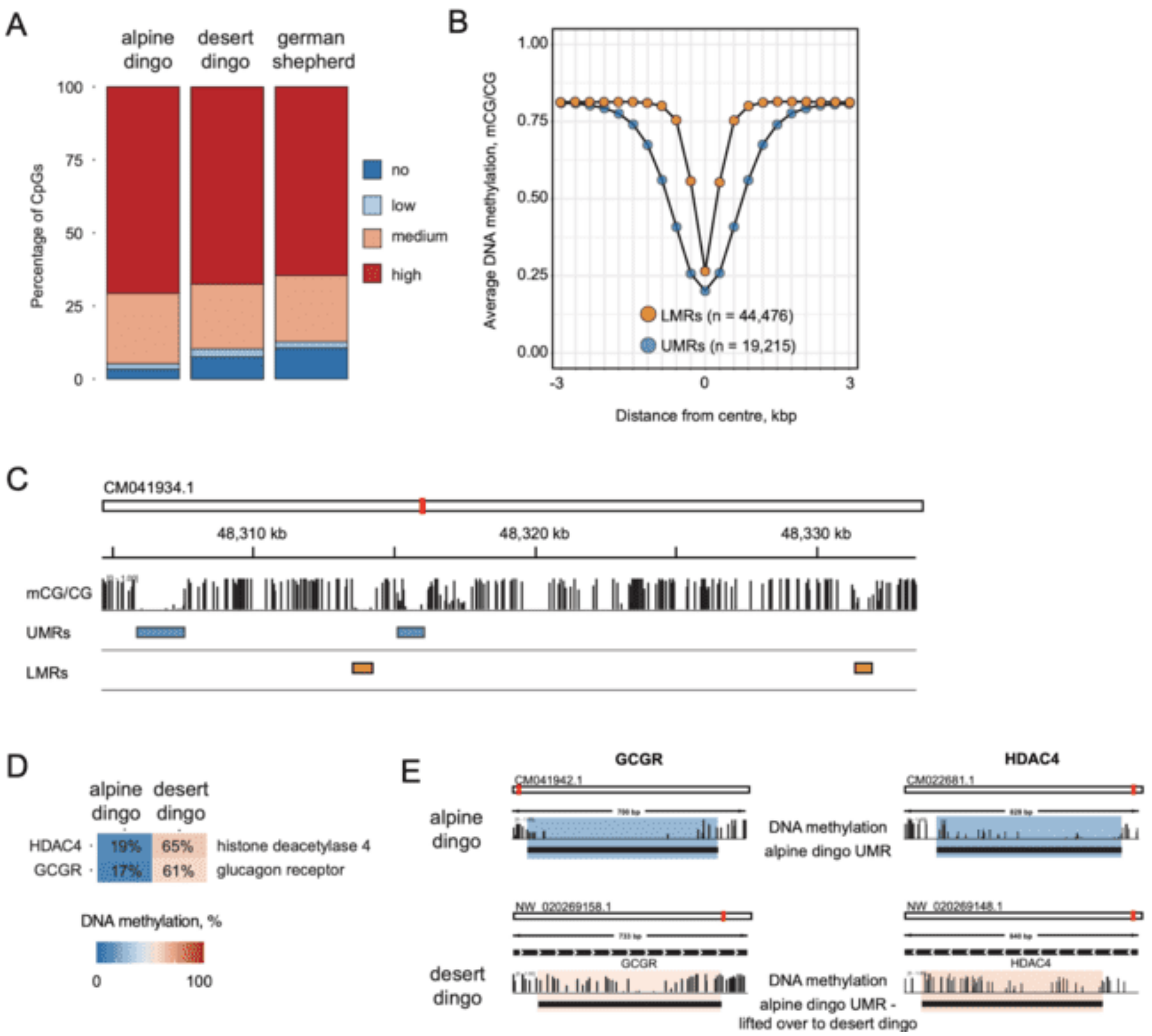


**Supplementary Figure 8:** DNA methylation profiling of alpine dingo Cooinda's whole blood

**Supplementary Figure 8 Legend:** (A) Percentage of CpG sites with different levels of methylation. High, 80-100%; medium, 20-80%; low, >0-20%; no, 0%. **(B)** Average DNA methylation profiles of hypomethylated regions into CpG-rich unmethylated regions (UMRs) and CpG-poor low-methylated regions (LMRs). **(C)** Integrative Genomics Viewer **(**IGV) browser track depicting DNA methylation profile and putative regulatory elements (UMRs and LMRs). **(D)** Heatmap depicting average DNA methylation at hypomethylated UMRs in the alpine dingo genome, which are more than 50% methylated in the desert dingo genome. **(E)** IGV browser track depicting hypomethylated UMRs within GCGR and HDAC4 genes in the alpine dingo genome, which are hypermethylated in the desert dingo genome.

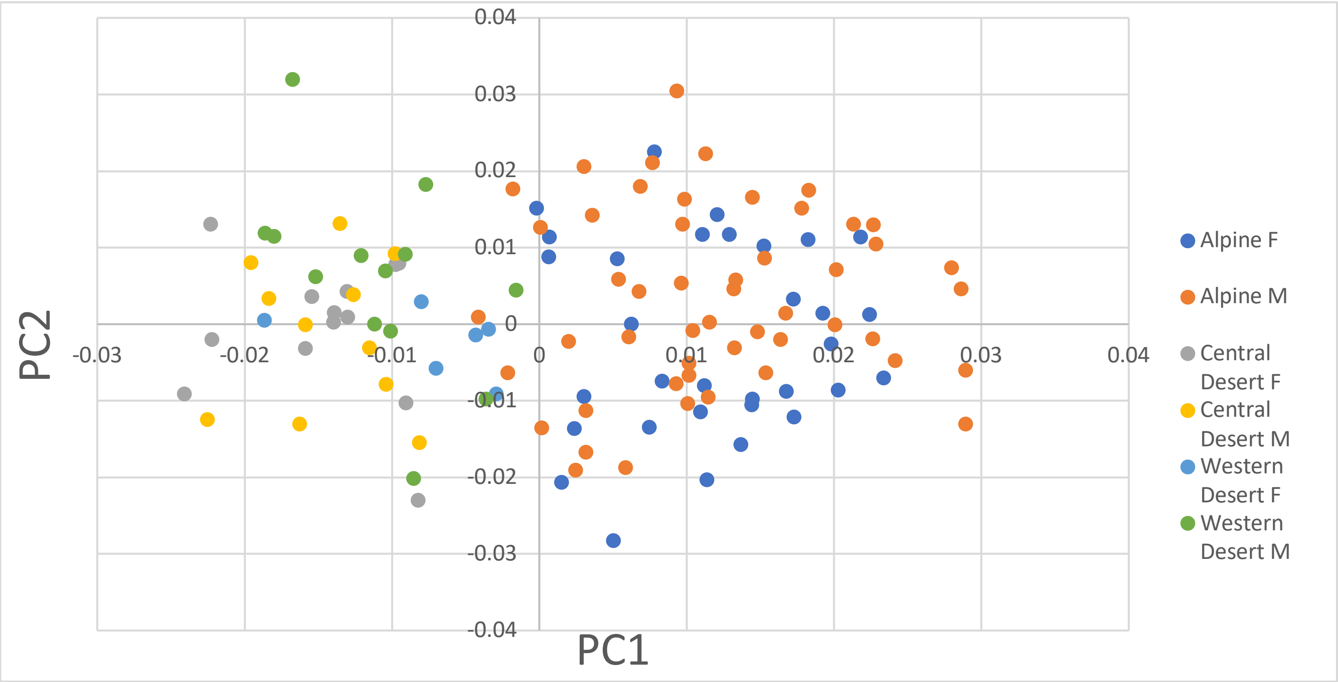


**Supplementary Figure 9**

**Supplementary Figure 9 Legend:** Scatterplot of PC1 and PC2 values for sexed dingo specimens. The distribution of greater PC2 values slightly favors males in all populations except for Central Desert, which is a very gracile population with relatively minimal differences between the cranial morphology of different sexes. In general, however, the difference in PC2 between males and females in any population is very marginal and neither greater nor lesser values are particularly strongly associated with either sex.

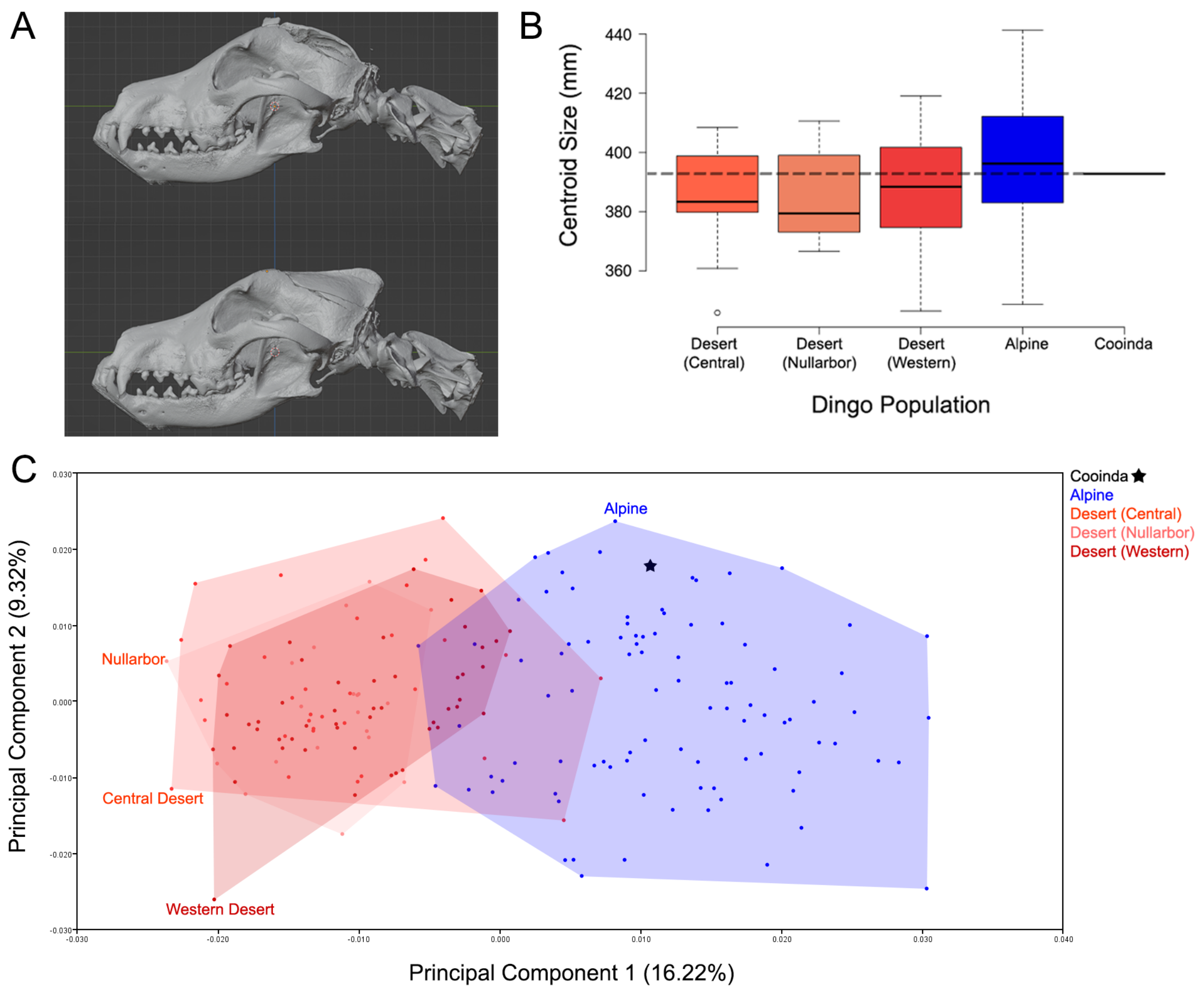


**Supplementary Figure 10:** Cooinda's cranial morphology

**Supplementary Figure 10 Legend:** (A) Cranium before (upper) and after (lower) cranial reconstruction (lower). This was required because the brain was removed immediately after death, which caused some damage to the braincase. (B) Cranium size. Cooinda's cranium is larger than the median size reported for Desert dingoes, in line with Alpine dingoes in general, although this difference is not major and there is heavy overlap between the two regions. Her centroid size (392.80mm) is slightly below the pooled Alpine mean (396.49mm) and median (396.23mm), but well below the mean (403.26mm) and median (403.64mm) for Alpine males specifically, which make up a majority of the sample (male $n$ = 50; female $n$ = 33; sex unknown $n$ = 9). Alpine dingoes, as with all regional dingo populations, exhibit significant sexual dimorphism in centroid size with males being on average 4.20% larger

[65]. (C) Principal component ordination of allometric residuals. The residuals of a regression of shape against log centroid size were plotted to further explore the role of size (allometry) in overall form. This revealed that the separation of Alpine and Desert populations, and Cooinda's position within the former, remains essentially identical to their original distribution s (Fig. 5a) when the size-related allometric component of form is removed from consideration.

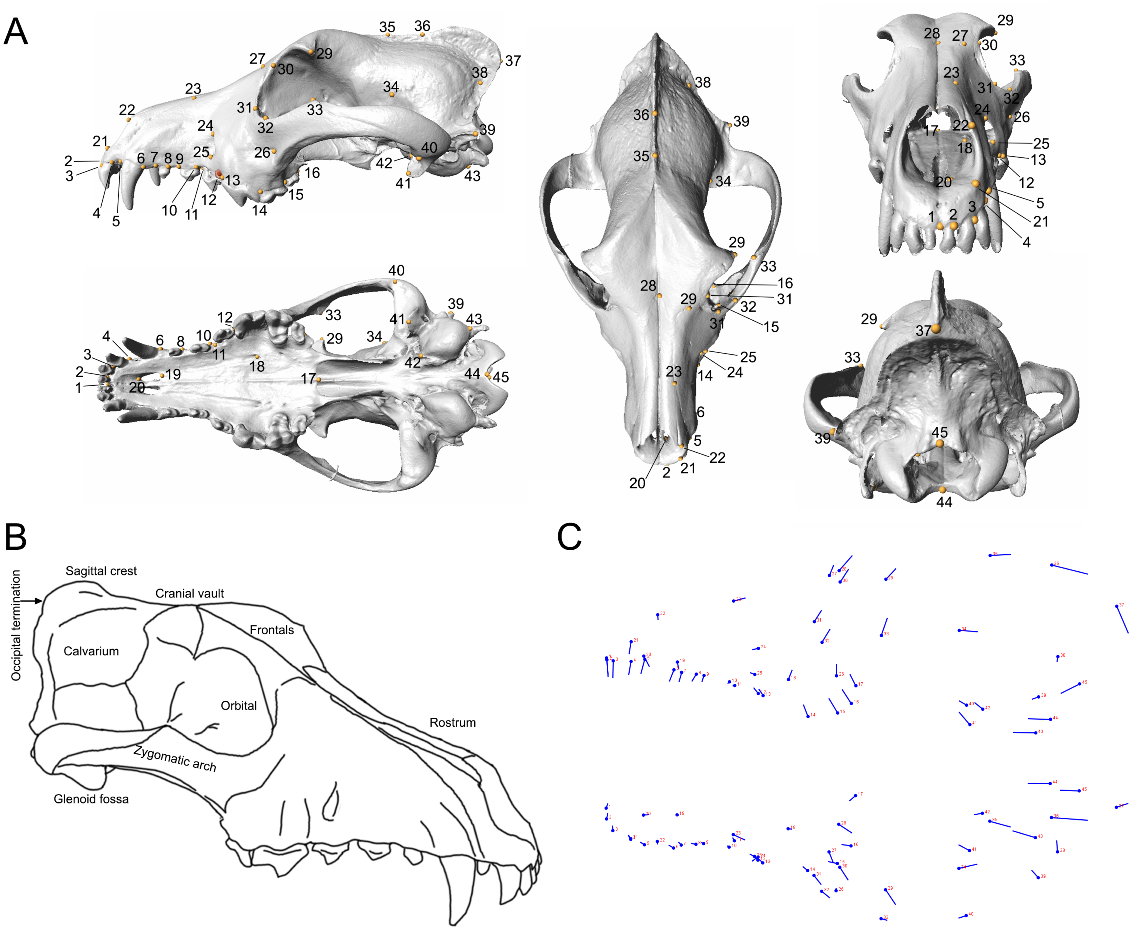


**Supplementary Figure 11**

**Supplementary Figure 11 Legend:** (A) Landmarks used in this study. (B) diagram of canid skull with basic anatomical features and regions referred to in-text. (C) Lollipop figures illustrating change in landmark positions along PC2 in lateral (upper) and dorsal (lower) views. The lollipop "head" represents the mean position, and the end of the "stick" represents its position with the highest PC1 score.

**Supplementary Table 1**. Alpine dingo SNVs and SVs summary by chromosome.

| **Chromosome** | **Alpine dingo SNV count vs** | | **Alpine dingo SV count vs** | | **Chromosome bp** |
|---|---|---|---|---|---|
| | **Desert dingo** | **CanFam4.0** | **Desert dingo** | **CanFam4.0** | |
| **1** | 77926 | 195865 | 1370 | 3086 | 123706252 |
| **2** | 55068 | 134823 | 1192 | 2074 | 85446638 |
| **3** | 63384 | 162076 | 939 | 2552 | 92885137 |
| **4** | 53989 | 145400 | 1001 | 2287 | 88750986 |
| **5** | 57509 | 137691 | 981 | 2181 | 89373516 |
| **6** | 49987 | 114964 | 1067 | 2510 | 77892171 |
| **7** | 61659 | 120334 | 1132 | 2074 | 82078758 |
| **8** | 51280 | 132310 | 1038 | 2138 | 74537974 |
| **9** | 48343 | 87955 | 1027 | 1566 | 63158580 |
| **10** | 42060 | 106101 | 754 | 1477 | 71288900 |
| **11** | 54954 | 119566 | 1100 | 1950 | 75713417 |
| **12** | 80643 | 119102 | 1661 | 2123 | 73055177 |
| **13** | 67246 | 96899 | 1076 | 1411 | 65408697 |
| **14** | 44691 | 97750 | 899 | 1913 | 61550799 |
| **15** | 56205 | 93452 | 1023 | 1627 | 65319253 |
| **16** | 51884 | 99759 | 1176 | 2021 | 62499104 |
| **17** | 35216 | 114685 | 754 | 1783 | 64752584 |
| **18** | 50255 | 93879 | 1160 | 1536 | 56713732 |
| **19** | 54419 | 94065 | 988 | 1311 | 55746904 |
| **20** | 40988 | 86977 | 703 | 1446 | 58849343 |
| **21** | 41147 | 96550 | 691 | 1293 | 51792689 |
| **22** | 33737 | 94950 | 636 | 1604 | 61776113 |
| **23** | 44240 | 87440 | 794 | 1274 | 53490029 |
| **24** | 32824 | 81112 | 491 | 1037 | 48051091 |
| **25** | 43100 | 97505 | 739 | 1413 | 51754766 |
| **26** | 38604 | 91768 | 757 | 1406 | 40258947 |
| **27** | 52772 | 76375 | 1075 | 1447 | 46564242 |
| **28** | 29616 | 81846 | 517 | 984 | 41881905 |
| **29** | 29642 | 75460 | 507 | 1121 | 42159209 |
| **30** | 22298 | 69332 | 486 | 1074 | 40431870 |
| **31** | 41002 | 82966 | 628 | 1015 | 40406310 |

| | | | | | |
|---|---|---|---|---|---|
| **32** | 26587 | 83791 | 614 | 1610 | 39322259 |
| **33** | 33909 | 57901 | 692 | 977 | 31731374 |
| **34** | 35988 | 89665 | 567 | 1843 | 42557588 |
| **35** | 24494 | 56238 | 383 | 683 | 26863133 |
| **36** | 19944 | 53165 | 393 | 796 | 31103624 |
| **37** | 31481 | 49270 | 789 | 935 | 31168689 |
| **38** | 26115 | 58654 | 416 | 780 | 25215458 |
| **X** | 24584 | 102071 | 582 | 2166 | 125292608 |

**Supplementary Table 2.** Distance matrix table showing SNVs above diagonal and Indels below. All possible pairwise alignments were generated using MUMmer4 [44] (v4.0.0 beta 2) and SNVs/indels numbers calculated using MUMmer4 ‘show-snp’ script.

| | **Desert** | **Alpine** | **Basenji1 (China)** | **Basenji2 (Wags)** | **GSD1 (Nala)** | **GSD2 (Mischa)** | **Labrador** | **Boxer** | **Great Dane** | **Greenland Wolf** |
|---|---|---|---|---|---|---|---|---|---|---|
| **Desert** | - | 1934204 | 4379273 | 4058304 | 4157347 | 4099899 | 4266975 | 3956320 | 3858069 | 5039138 |
| **Alpine** | 3525802 | - | 4351866 | 4048746 | 4125867 | 4061800 | 4219881 | 3939675 | 3802100 | 4696525 |
| **Basenji1** | 6813866 | 6946862 | - | 2199194 | 3893739 | 3855744 | 3922731 | 3700555 | 3605316 | 5155027 |
| **Basenji2** | 6482039 | 6567647 | 4372616 | - | 3515180 | 3471928 | 3686649 | 3375257 | 3246911 | 4949577 |
| **GSD1** | 6290364 | 6362924 | 6237235 | 5742582 | - | 2064119 | 3477794 | 3101130 | 3007546 | 5078984 |
| **GSD2** | 6229282 | 6301226 | 6186934 | 5663021 | 3396958 | - | 3426212 | 3072348 | 2990169 | 5049776 |
| **Labrador** | 7072684 | 7122926 | 6758598 | 6529100 | 6029553 | 5975640 | - | 3174798 | 3162230 | 5252186 |
| **Boxer** | 6124985 | 6229361 | 6004983 | 5678411 | 5061694 | 5025968 | 5816010 | - | 2773431 | 4947075 |
| **Great Dane** | 6601081 | 6693070 | 6455860 | 6160610 | 5582788 | 5578574 | 6440463 | 5255792 | - | 4776954 |
| **Greenland Wolf** | 7273469 | 6925872 | 7606227 | 7343748 | 7339762 | 7266186 | 8130863 | 7190906 | 7642878 | - |

GSD is German shepherd dog

**Supplementary Table 3.** Mean and median PC2 scores for different sexes from dingo populations
.

| | PC2 Mean | PC2 Median |
|---|---|---|
| Alpine F | -0.00203 | -0.00481572 |
| Alpine M | 0.003143 | 0.001463463 |
| Central Desert F | -0.00062 | 0.000931911 |
| Central Desert M | -0.00131 | -3.97883E-05 |
| Western Desert F | -0.00226 | -0.001064826 |
| Western Desert M | 0.00602 | 0.006917365 |

**Supplementary Table 4.** List and description of cranial landmarks used in this study. After Koungoulos [65].

| *Landmark #* | *Description* |
|---|---|
| *1* | *Between left and right first incisors.* |
| *2* | *Between first and second incisor.* |
| *3* | *Between second and third incisor.* |
| *4* | *Posterior corner of third incisor alveolus.* |
| *5* | *Anterior corner of canine alveolus.* |
| *6* | *Posterior corner of canine alveolus.* |
| *7* | *Anterior corner of first premolar alveolus.* |
| *8* | *Posterior corner of first premolar alveolus.* |
| *9* | *Anterior corner of second premolar alveolus.* |
| *10* | *Posterior corner of second premolar alveolus.* |
| *11* | *Anterior corner of third premolar alveolus.* |
| *12* | *Posterior corner of third premolar alveolus.* |
| *13* | *Anterior corner of fourth premolar (carnassial) alveolus.* |
| *14* | *Posterior corner of fourth premolar (carnassial)/Anterior corner of first molar alveoli.* |
| *15* | *Posterior corner of first molar (carnassial)/Anterior corner of second molar alveoli.* |
| *16* | *Posterior edge of second molar alveolus.* |
| *17* | *Staphylion at edge of palate and choanal region.* |
| *18* | *Greater palatal foramen.* |
| *19* | *Proximal end of palatine fissure.* |
| *20* | *Distal end of palatine fissure.* |
| *21* | *Lower corner of distal incisive bone.* |
| *22* | *Upper corner of distal incisive bone.* |
| *23* | *Intersection of incisive, maxilla and nasal bones.* |
| *24* | *Upper end of infraorbital foramen ridge.* |
| *25* | *Lower end of infraorbital foramen ridge.* |
| *26* | *Lower limit of intersection of maxilla and zygomatic bones.* |
| *27* | *Upper limit of intersection of maxilla and frontal bones.* |
| *28* | *Upper limit of intersection of nasal and frontal bone.* |
| *29* | *Furthest extent of zygomatic process of frontal bone.* |
| *30* | *First "corner" of orbital rim.* |
| *31* | *Second "corner" of orbital rim.* |
| *32* | *Third "corner" of orbital rim.* |
| *33* | *Fourth "corner" of orbital rim.* |
| *34* | *Intersection of frontal, parietal and temporal bones.* |
| *35* | *Intersection of cranial midline with frontal-temporal boundary suture (bregma).* |
| *36* | *Intersection of the parietal and occipital bones on the sagittal crest.* |
| *37* | *Most proximal extent/tip of occipital protuberance (occiput).* |
| *38* | *Intersection of nuchal crest with parietal-temporal boundary suture.* |
| *39* | *Lower flare of nuchal crest.* |
| *40* | *Underside of mandibular fossa, where zygomatic process begins.* |

| | |
|---|---|
| *41* | *Underside of retroarticular process.* |
| *42* | *Anterior corner of auditory bulla.* |
| *43* | *Posterior corner of auditory bulla.* |
| *44* | *Lower central rim of foramen magnum, between occipital condyles.* |
| *45* | *Upper central rim of foramen magnum, between occipital condyles.* |
